# Deleting a UBE3A substrate rescues impaired hippocampal physiology and learning in Angelman syndrome mice

**DOI:** 10.1101/625418

**Authors:** Gabrielle L. Sell, Wendy Xin, Emily K. Cook, Mark A. Zbinden, Thomas B. Schaffer, Robert N. O’Meally, Robert N. Cole, Antonello Bonci, Seth S. Margolis

## Abstract

In humans, loss-of-function mutations in the *UBE3A* gene lead to the neurodevelopmental disorder Angelman syndrome (AS). AS patients have severe impairments in speech, learning and memory, and motor coordination, for which there is currently no treatment. In addition, *UBE3A* is duplicated in >1-2% of patients with autism spectrum disorders – a further indication of the significant role it plays in brain development. Altered expression of UBE3A, an E3 ubiquitin ligase, is hypothesized to lead to impaired levels of its target proteins, but identifying the contribution of individual UBE3A targets to UBE3A-dependent deficits remains of critical importance. Ephexin5 is a putative UBE3A substrate that has restricted expression early in development, regulates synapse formation during hippocampal development, and is abnormally elevated in AS mice, modeled by maternally-derived *Ube3a* gene deletion. Here, we report that Ephexin5 is a direct substrate of UBE3A ubiquitin ligase activity. Furthermore, removing Ephexin5 from AS mice specifically rescued hippocampus-dependent behaviors, CA1 physiology, and deficits in dendritic spine number. Our findings identify Ephexin5 as a key driver of hippocampal dysfunction and related behavioral deficits in AS mouse models. These results demonstrate the exciting potential of targeting Ephexin5, and possibly other UBE3A substrates, to improve symptoms of AS and other UBE3A-related developmental disorders.

## INTRODUCTION

Angelman syndrome (AS) is a neurodevelopmental disorder that affects approximately one in 15,000 individuals and is characterized by severe motor dysfunction, loss of speech, frequent seizures, and debilitating cognitive impairments (Bird, 2014; Williams, 2005). The majority of AS patients carry genomic lesions that disrupt the E3 ubiquitin ligase UBE3A (Kishino et al., 1997). In humans, the *UBE3A* gene resides within chromosomal region 15q11-13 and is subject to brain-specific genomic imprinting, with predominant transcription of the maternal allele in neuronal cells across the brain (Albrecht et al., 1997; Clayton-Smith and Laan, 2003). Inheritance of an abnormal, inactive maternal copy of the *UBE3A* gene is currently thought to account for 85-90% of AS cases (Clayton-Smith and Laan, 2003). Consistent with this, deletion of the maternal copy of UBE3A in mice (AS mice) leads to clear cellular, electrophysiological, and behavioral deficits in these animals (Jiang et al., 1998).

*UBE3A* encodes a HECT (Homologous to the E6-AP Carboxyl Terminus) domain E3 ubiquitin ligase that catalyzes the addition of ubiquitin to lysine residues on substrate proteins. Several mutations identified in AS patients specifically disrupt this ubiquitin ligase activity of UBE3A while still expressing full-length protein (Cooper et al., 2004). UBE3A-dependent ubiquitylation is primarily thought to target substrates for degradation by the 26S proteasome (Matentzoglu and Scheffner, 2008). Ubiquitin-dependent proteasome degradation clears unwanted proteins and is necessary to maintain proper intracellular protein homeostasis and neuronal function (Yi and Ehlers, 2005). Thus, the possibility that aberrant levels of UBE3A targets are causing the numerous phenotypic deficits observed in AS is a prevailing hypothesis in the AS field. This hypothesis has generated excitement among AS researchers as it could lead to the development of therapeutic strategies targeting specific UBE3A substrates. Currently, however, the involvement of direct UBE3A substrates in AS phenotypes remains largely untested.

UBE3A is highly expressed in the hippocampus, a brain region that is critical for learning and memory (Burette et al., 2017). Ephexin5, a RhoA guanine nucleotide exchange factor, is a candidate UBE3A substrate that is also enriched in the hippocampus. During development, Ephexin5 acts as a brake on excitatory synapse formation (Margolis et al., 2010). Previous work identified Ephexin5 as a probable substrate of the ubiquitin-proteasome system and reported a decrease in total ubiquitylated Ephexin5 in AS mice (Margolis et al., 2010). Based on these data, we hypothesized that Ephexin5 is a direct substrate of UBE3A and that in AS mice, increased expression of Ephexin5 would be critically involved in UBE3A-dependent hippocampal dysfunction. Here we report that Ephexin5 is a direct substrate of UBE3A and is elevated in adult AS mice brains. Importantly, removal of Ephexin5 rescued hippocampus-related impairments in behavior, spine density, and electrophysiology in the CA1 region of AS mice. These data demonstrate the role of a UBE3A substrate in AS cognitive pathology and pinpoints Ephexin5 as a promising target for treating cognitive deficits in AS.

## MATERIALS AND METHODS

(See Table 1 for detailed key resources).

**Table 1:** A detailed outline of critical reagents and resources required for this study.

| REAGENT or RESOURCE | SOURCE | IDENTIFIER |
| --- | --- | --- |
| **Antibodies** |  |  |
| Mouse anti-UBE3A (330) | Sigma-Aldrich | Cat# E8655; RRID AB_261956 |
| Mouse anti-UBE3A (3E5) | Sigma-Aldrich | Cat# SAB1404508; RRID AB_10740376 |
| Mouse anti-β-Actin (8226) | Abcam | Cat# ab8226; RRID AB_306371 |
| Chicken anti-GFP | Aves Labs | Cat#1020; RRID AB_10000240 |
| Rabbit anti-Ephexin5 | M. Greenberg (Harvard) | (Margolis et al., 2010) |
| Goat anti-Rabbit IgG, (H+L), HRP | Cell Signaling Technology | Cat# 7074S; RRID AB_2099233 |
| Horse anti-Mouse IgG, (H+L), HRP | Cell Signaling Technology | Cat# 7076S; RRID AB_330924 |
| Donkey anti-Mouse Cy3 | Jackson ImmunoResearch | Cat# 715-165-151; RRID AB_2315777 |
| Goat anti-Chicken 488 | Thermo Fisher Scientific | Cat# A-11039; RRID AB_142924 |
| **Bacterial and Virus Strains** |  |  |
| DH5α | Thermo Fisher Scientific | Cat# 18265017 |
| Rosetta DE3 BL21 bacteria | E. Goley (kind gift) | N/A |
| **Chemicals, Peptides, and Recombinant Proteins** |  |  |
| Poly-L-Lysine (PLL) | Sigma-Aldrich | Cat# P4707 |
| Papain | Worthington | Cat# LS003127 |
| B27 | Thermo Fisher Scientific | Cat# 17504044 |
| Neurobasal | Thermo Fisher Scientific | Cat# 21103049 |
| Neurobasal A | Thermo Fisher Scientific | Cat# 10888-022 |
| Penicillin/streptomycin | Sigma-Aldrich | Cat# 15140122 |
| L-Glutamine | Thermo Fisher Scientific | Cat# 25030081 |
| Trypsin | Thermo Fisher Scientific | Cat# 25200056 |
| Trypsin Inhibitor | Sigma-Aldrich | Cat# T9253; CAS: 9035-81-8 |
| Trypsin/LysC (MS/MS) | Promega | Cat# V5071 |
| DMEM | Thermo Fisher Scientific | Cat# 11960069 |
| Fetal bovine serum | Thermo Fisher Scientific | Cat# 16000044 |
| Phosphate buffered saline | Thermo Fisher Scientific | Cat# 10010049 |
| cOmplete Protease inhibitor | Sigma-Aldrich | Cat# 11836170001 |
| Precision Plus Standards | Bio-Rad | Cat# 1610374 |
| PROTOGEL (30%) | Thermo Fisher Scientific | Cat# 50-899-90118 |
| Lipofectamine 2000 | Life Technologies | Cat# 11668019 |
| UBE1 | Boston Biochem | Cat# E-304 |
| E6AP | Boston Biochem | Cat# E3-230 |
| E6 | Boston Biochem | Cat# AP-120 |
| Ubiquitin | Boston Biochem | Cat# U-100H |
| Mg-ATP | Boston Biochem | Cat# B-20 |
| TCEP | Thermo Fisher Scientific | Cat# 20490 |
| Triton-X 100 | Sigma-Aldrich | Cat# T9284 |
| Paraformaldehyde | Thermo Fisher Scientific | Cat# 50980487 |
| Neg50 OCT | Thermo Fisher Scientific | Cat# 6502 |
| Fluoromount-G | Southern Biotech | Cat# 0100-01 |
| Hoechst | Molecular Probes | Cat# H-3570 |
| DNaseI | Qiagen | Cat# 79254 |
| DMSO | Thermo Fisher Scientific | Cat# 25-950-CQC |
| lysozyme | Sigma-Aldrich | Cat# L6876 |
| Clozapine *N*-oxide (CNO) | Tocris | Cat# 4936; CAS: 34233-69-7 |
| TTX | Tocris Bioscience | Cat# 1078; CAS: 4368-28-9 |
| Gabazine | Sigma-Aldrich | Cat# SR-95531; CAS: 104104-50-9 |
| APV | Tocris Bioscience | Cat# S106; CAS: 79055-68-8 |
| 2-mercaptoethanol | Bio-Rad | Cat# 1610710 |
| Nitric Acid | Thermo Fisher Scientific | Cat# A200212 |
| NaCl | Sigma-Aldrich | Cat# S9625 |
| KCl | Sigma-Aldrich | Cat# P9541 |
| IPTG | Sigma-Aldrich | Cat# I5502 |
| Tris base | Sigma-Aldrich | Cat# T1503 |
| PMSF | Sigma-Aldrich | Cat# P7626; CAS: 329-98-6 |
| DTT | Sigma-Aldrich | Cat# D5545 |
| LB broth | Sigma-Aldrich | Cat# L3022 |
| LB plates | Quality Biologicals | Cat# 340-107-231 |
| Critical Commercial Assays |  |  |
| PureLink HiPure Plasmid Filter Maxiprep Kit | Invitrogen | Cat# K210016 |
| Experimental Models: Organisms/Strains |  |  |
| C57Bl/6j | Jackson Laboratories | RRID:IMSR_JAX:000664 |
| Ephexin5<sup>-/-</sup> 129 | M. Greenberg (Harvard) | (Margolis et al., 2010) |
| Ephexin5<sup>-/-</sup> C57BL/6 | This study | (Margolis et al., 2010) |
| 129-*Ube3a*<sup>tm1Alb</sup>/J | Jackson Laboratories | RRID:IMSR_JAX:004477 |
| B6-*Ube3a*<sup>tm2Alb</sup>/J<br>(backcrossed – this study) | Jackson Laboratories | RRID:IMSR_JAX:017765 |
| tdTomato cre reporter lines | Jackson Laboratories | RRID:IMSR_JAX:007909 |
| CamK2a-cre | Jackson Laboratories | RRID:IMSR_JAX:005359 |
| Tg(Thy1-EGFP) MJrs/J | Jackson Laboratories | RRID:IMSR_JAX:007788 |
| Floxed-UBE3A | B. Philpot (kind gift) | (Judson et al., 2016) |
| Recombinant DNA |  |  |
| ube3a-shRNA | M. Greenberg (Harvard) | (Greer et al., 2010) |
| scr-shRNA | M. Greenberg (Harvard) | (Greer et al., 2010) |
| pcDNA5/FRT-HA-hM4D(Gi) | (Armbruster et al., 2007) | Addgene Plasmid # 45548 |
| EGFP | M. Greenberg (Harvard) | (Margolis et al., 2010) |
| pGEX-GST-Ephexin5 - WT | This Study | N/A |
| Software and Algorithms |  |  |
| Proteome Discoverer (v1.4) | Thermo Fisher Scientific | Cat# OPTON-30795 |
| Mascot (v2.5.1) | Matrix Science | www.matrixscience.com |
| Scaffold (v4.8.4) | Proteome Software | http://www.proteomesoftware.com/products/scaffold/ |
| Fiji/ImageJ | National Institutes of Health | https://imagej.nih.gov/ij/docs/guide/146-2.html |
| Metamorph | Molecular Devices |  |
| NeuronStudio | (Rodriguez et al., 2008) | http://research.mssm.edu/cnic/tools-ns.html |
| MiniAnalysis program (v 6.0) | Synaptosoft | http://www.synaptosoft.com/MiniAnalysis/ |
| Other |  |  |
| 12 mm Glass coverslips | Bellco | Cat# 1943-10012A |
| Glutathione Sepharose | GE Healthcare | Cat # 17513201 |
| Slide-A-Lyzer | Thermo Fisher Scientific | Cat # 66380 |

### Mice

All animals were socially housed, with food *ad libitum* on a 14:10 hour light/dark cycle. Ephexin5^+/-^ males were crossed with Ube3A^m+/p-^/Ephexin5^+/-^ females to obtain animals that were Ube3A wild type or maternally-deficient and Ephexin5 wild type or null. These crosses were performed in isogenic strains for both 129S7 and C57Bl/6J that had been backcrossed 10 generations, given previous published evidence that a single genetic background in these animals may not be sufficient for determining all phenotypes (Sittig et al., 2016). To label neurons fluorescently for spine analysis, 129S7 Ube3A^m+/p-^/Ephexin5^+/-^ were crossed with Thy1-EGFP mice (Jackson STOCK Tg(Thy1-EGFP)MJrs/J, 007788), a C57Bl/6j strain, allowing the use of an F1 generation as previously published (Copping et al., 2017; Sittig et al., 2016). From this cross, Ephexin5^+/-^ males were crossed with Ube3A^m+/p-^/Ephexin5^+/-^ females, with the EGFP present on either the male or female. WT, AS, AS/E5^-/-^, and E5^-/-^ animals positive for EGFP were used for analysis. C57Bl/6j CamKIIa-Cre mice were obtained, and expression pattern of this line was determined by crossing these animals to a TdTomato reporter line (Cre^+^/TdTomato^+^ animals, Fig. 2A). To selectively remove Ube3A expression from excitatory forebrain neurons, the CamKIIa-Cre line was crossed to floxed-Ube3A mice (Judson et al., 2016). Ube3A expression was determined by Western blot analysis after experiments (Fig. 2B). Both sexes were used for all experiments and no sex-specific alterations were discovered. The health of the Ephexin5-null animals has been previously published and we, similarly, saw no overt health issues (Margolis et al., 2010; Sell et al., 2017).

### Neuronal cultures

For primary mouse neuronal cultures, wild-type C57BL/6 mice or C57BL/6 Ephexin5^-/-^ mice were sacrificed at P0 and dissected. Brains were removed and dissected cortices were then moved into dissociation buffer (0.3% BSA, 12 mM MgSO_4_, 10 mM HEPES, 0.6% glucose in Hank’s Balanced Salt Solution) plus 16.67 U/ml Papain, which was incubated for 30 minutes at 37 °C. All neurons were mixed together and contain both male and female neuronal tissue. It was challenging to tell the sex of a mouse embryo and there was not enough material to separate out each embryo and still be able to perform the described biochemical studies in this manuscript. Authenticity of neuronal cultures was in part determined by morphology using light microscopy and immunoblotting of lysates from neuronal cultures with well-validated antibodies to neuronal enriched markers. Proteolyzed tissue was rinsed for 5 minutes twice in 10 mg/ml Trypsin inhibitor. Tissue was then resuspended in Neurobasal and mechanically dissociated into a single-cell suspension. Cells were counted with a hemocytometer and neurons were plated for imaging on glass coverslips within wells of a 24-well plate at a concentration of 200,000 cells/well.

### DNA constructs

The following plasmid constructs were gifts from Bryan Roth: pcDNA5/FRT-HA-hM4D(Gi) (Addgene: 45548) (Armbruster et al., 2007). UBE3A shRNA and UBE3A scrambled shRNA (sc-shRNA) were previously described (Greer et al., 2010).

### Antibodies

Ephexin5 antibodies, raised in rabbit against a GST-fusion protein containing Ephexin5 amino acids 1-418, were previously described (Margolis et al., 2010). The following antibodies are commercially available and used according to manufacturer’s suggestions for immunoblotting immunofluorescence studies: UBE3A (Sigma-Aldrich E8655) for Western, UBE3A (Sigma-Aldrich 3E5) for IF, β-Actin (Abcam ab8226), GFP (Aves 1020). Secondary antibodies are described below.

### Immunoblotting

For immunoblot analysis, antibodies to Ephexin5 (1:1000, 5% non-fat milk, TBS, 0.05% Tween 20), Actin (1:5000, 5% BSA, TBS, 0.05% Tween 20), and UBE3A (1:2000, 5% non-fat milk, TBS, 0.05% Tween 20) were utilized. Anti-mouse IgG and anti-rabbit IgG secondary antibodies (Cell Signaling Technology 7076S and 7074S, respectively) were used for immunoblotting at a concentration of 1:5000 in the same blocking buffer as the primary antibody. Optical density of the immunoblot bands was quantified using ImageJ.

### Expression and purification of GST-Ephexin5

For the generation of purified full-length Ephexin5 protein, a pGEX plasmid encoding N-terminally fused GST-Ephexin5 was transformed into Rosetta DE3 BL21 bacteria. Starter cultures harboring Ephexin5 plasmid were grown overnight at 37 °C and used to inoculate larger cultures the following day at 1:100. Cultures were grown at 37 °C until reaching an OD (600) of 0.6-0.8, at which point cultures were divided into 100 mL volumes and grown at 16 °C for 1 hour. GST-Ephexin5 expression was induced by addition of 0.5 μM Isopropyl β-D-1-thiogalactopyranoside and cultures were grown overnight at 16 °C. Cells were spun and harvested in 2.3 M sucrose, 50 mM Tris pH 7.5, and 1 mM EDTA. Recovered cells were lysed on ice for 1 hour in 50 mM Tris pH 7.5, 10 mM KCl, 1 mM EDTA, 2 mM DTT, 1 mM PMSF, and 55 mM lysozyme. Lysates were then treated with 0.1% sodium deoxycholate, 25 mM MgCl_2_, 0.4 nM DNase I for 15 minutes and insoluble material was separated from lysate via centrifugation at 4,000 x g for 30 minutes at 4 °C. To the supernatant, comprised of bacterial lysate containing overexpressed GST-Ephexin5, Glutathione Sepharose beads were added and incubated under constant rotation at 4 °C for 2-4 hours. Beads with bound GST-Ephexin5 protein were then rinsed 4 times with 10 mM HEPES pH 7.5, 1 mM DTT, 300 mM NaCl and once with 10 mM HEPES pH 7.5, 1 mM DTT. Protein was then eluted twice from the beads with 50mM Tris pH8, 10mM reduced glutathione. Elutions were dialyzed into 50 mM HEPES pH 8.0, 200 mM NaCl, 10% glycerol, 1 mM TCEP using Slide-A-Lyzer Dialysis Cassette (Extra Strength) with a 10,000 MW cutoff. Dialyzed protein was aliquoted and stored at −80°C and aliquots were taken for *in vitro* ubiquitylation assays.

### In Vitro Ubiquitination Reaction

The UBE3A in vitro ubiquitination reactions were prepared with the following components from Boston Biochem: 3 μl 10X reaction buffer (500 mM HEPES, 500 mM NaCl, 10 mM TCEP, pH 8), 50 nM UBE1, 1uM UBE2L3, 1 μM E6AP, 1 μM E6, 50 μM Ubiquitin, 10mM Mg-ATP, and ∼0.79 ug GST-Ephexin5. Reactions were prepared according to manufacturer’s instructions. Briefly, reactions were incubated at 37 °C for 90 or 180 min. Reactions were stopped using Lammeli Buffer to be run on SDS-PAGE for western blotting or flash frozen to be used for mass spectrometry analysis.

### Mass Spec ID of Ubiquitylations Sites

Following *in vitro* UBE3A ubiquitylation, Ephexin5 was reduced 5 μM dithiothreitol (DTT) 20 mM ammonium bicarbonate at pH 8.5 at 60 °C for 1 hr, and after cooling, alkylated with 10 μM iodoacetomide for 15 minutes at room temperature in the dark. Reduced and alkylated proteins were digested overnight at 37 °C by adding 1:20 Trypsin/LysC mixture: protein in 20 mM ammonium bicarbonate. Peptides from protein digests were desalted using Oasis HLB uElution solid phase extraction plates (Waters) equilibrated and washed with 0.1% TFA, then eluted with 60% acetonitrile in 0.1% TFA and dried by vacuum centrifugation. Desalted peptides were resuspended in 10 μl 2% acetonitrile in 0.1% formic acid and analyzed by reverse phase liquid chromatography coupled to tandem mass spectrometry. Peptides were separated on a picofrit house packed 75 μm x 200 mm ProntoSIL-120-5-C18 H column (5 µm, 120 Å (BISCHOFF), www.bischoff-chrom.com) using 2-90% acetonitrile gradient at 300 nl/min over 90 min on a EasyLC nanoLC 1000 (Thermo Scentific). Eluting peptides were sprayed through 10 µm emitter tip (New Objective, www.newobjective.com) at 2.2 kV directly into a Orbitrap Fusion (Thermo Scientific) mass spectrometer. Survey scans (full ms) were acquired from 350-1800 m/z with data dependent isolating the highest number of precursors in a 3 second cycle between each survey scan. Each peptide was isolated with a 1.6 Da window and fragmented with an HCD normalized collision energy of 30 and 15 s dynamic exclusion. Precursor and the fragment ions were analyzed at resolutions 120,000 and 30,000, respectively, with automatic gain control (AGC) target values at 4e5 with 50 ms maximum injection time (IT) and 1e5 with 100 ms maximum IT, respectively. Isotopically resolved masses in precursor (MS) and fragmentation (MS/MS) spectra were extracted from raw MS data without deconvolution and with deconvolution using MS2 Processor in Proteome Discoverer (PD) software. All extracted data were searched using Mascot against the RefSeq2017 protein database with mammalian taxonomy. The following criteria were set for all database searches: sample’s species; trypsin as the enzyme, allowing one missed cleavage, and the variable modifications of cysteine carbamidomethylation, Gly-Gly on lysine, methionine oxidation, asparagine and glutamine deamidation. Peptide identifications from Mascot searches were filtered at 1% False Discovery Rate (FDR) confidence threshold, based on a concatenated decoy database search, using Proteome Discoverer. Proteome Discoverer uses only the peptide identifications with the highest Mascot score for the same peptide matched spectrum from the different extraction methods. Mascot results were imported into Scaffold for validation and side-by-side comparisons of MS results.

### Behavioral Assays

All behavioral testing was done during the animal’s dark phase (without a reversed light/dark cycle) with littermate controls when the animals were 11-13 weeks old. The genotypes were blinded to the experimenter during both the tasks and analysis. Animals were habituated to testing room 20 minutes each day before tasks were begun. Any experiments performed included all 4 genotypes.

#### Novel Place Preference

Mice were habituated to the testing arena for 30 minutes in the testing room. The testing arena was a 10” x 10” box with a small visual cue on one side to orient the box. Between each mouse, the arena was cleaned with 70% ethanol. Twenty-four hours later, mice were exposed to 3 different objects for 10 minutes. An additional twenty-four hours later, mice were placed in the testing arena with the same 3 objects for 10 minutes, one object having been moved across the arena. Both days with the objects were video recorded. A blinded observer scored the amount of time each mouse spent investigating the objects using ODLog (Macropod Software). Investigation was defined as time spent with the mouse’s nose within 3 mm of the object, with orientation toward the object. Mice were excluded before unblinding if they did not investigate all objects. This was not restricted to any one genotype. C57Bl/6j WT (n=11), AS (n=13), AS/E5^-/-^ (n=10), E5^-/-^ (n=11). Object ((F=2, 123) = 24.56). Conditional *ube3A^loxp+^/camkIIa-cre^+^:* n=15, Kruskal-Wallis statistic = 2.498; *ube3A^loxp+^/camkIIa-cre*^-^: n=9, F(2, 24) = 0.3827; *ube3A^loxp+^/camkIIa-cre^+^*/E5^-/-^: n=7, F(2, 18) = 12.41; *ube3A^loxp+^/camkIIa-cre*^-^/E5^-/-^:n=12, Kruskal-Wallis statistic 11.07.

#### Novel Object Recognition

Similar to Novel Place Preference, except 129S7 mice were exposed to two identical objects. On the third day, instead of moving an object, one of the identical objects was replaced for a unique object. Analysis and timing of experiments is as described above. 129S7 WT (n=15), AS (n=12), AS/E5^-/-^ (n=18), E5^-/-^ (n=12). Genotype (df, t ratio): WT (28, 3.22096), AS (22, 1.17592), AS/E5^-/-^ (34, 2.8498), E5^-/-^ (22, 4.9558).

#### Passive Avoidance

C57Bl/6j and 129S7 mice were habituated to the chamber for 15 seconds before opening of the guillotine door separating the lit chamber from a dark chamber (Coulbourn Instruments). As mice entered the dark chamber, the door closed, mice were shocked (0.3 mA, 2 seconds) and immediately removed. 24 hours later, mice were placed in the lit box and allowed up to 5 minutes to enter the shock chamber. Latency on both days was recorded. Both chambers cleaned with 70% ethanol between mice. C57Bl/6j WT (n=5), AS (n=6), AS/E5^-/-^ (n=7), E5^-/-^ (n=7). Interaction (F (3,42) = 6.349). Day (F (1, 42) = 77.66). Genotype (F(3, 42) = 6.054). 129S7 WT (n=5), AS (n=4), AS/E5^-/-^ (n=4), E5^-/-^ (n=4). Interaction (F (3,26) = 0.5279). Day (F (1, 26) = 152.8). Genotype (F(3, 26) = 0.5279).

#### Rotarod

C57Bl/6j and 129S7 mice were habituated to the rotating rod for 10 minutes at a low speed. If animals fell off during this habituation, they were placed back on the rod until the 10 minutes was complete. After habituation, animals were placed on the rotating rod with increasing speed until the animal fell from the rod. The latency to fall was recorded. Each animal was put through 5 trials, with at least 10 minutes between each trial. For analysis, the highest and lowest latency for each animal was dropped, leaving 3 values per animal. The rotating rod was cleaned with 70% ethanol between each animal. C57Bl/6j WT (n=10), AS (n=11), AS/E5^-/-^ (n=4), E5^-/-^ (n=5). (F (3,26) = 14.71). 129S7 WT (n=11), AS (n=13), AS/E5^-/-^ (n=14), E5^-/-^ (n=10). (F (3, 44) = 6.956). Conditional *ube3A^loxp+^/camkIIa-cre*^+^: (n=16), *ube3A^loxp+^/camkIIa-cre*^-^: (n=8). t, df (0.6057, 22).

#### Marble Burying

16 marbles were evenly spaced in a 10”x10” arena with 2 inches of clean bedding. 129S7 mice were allowed 20 minutes to freely interact in the arena. Marbles that were completely covered, or covered at least 2/3 in bedding were counted as “buried”. Marbles were cleaned with 70% ethanol between each animal use. 129S7 WT (n=7), AS (n=8), AS/E5^-/-^ (n=7) and E5^-/-^ (n=6). (F (3, 24) = 10.16).

### Perfusion and Immunofluorescence

For *ex vivo* immunofluorescence analysis, 8 or 11 week old mice, as indicated, were perfused with ice cold 0.1 M phosphate buffer followed by 4% paraformaldehyde. Brains were removed, post-fixed in 4% paraformaldehyde and 30% sucrose for 4 days before being frozen in Neg50 OCT (Thermo Scientific) 30 micron sections were taken and stored in PBS with sodium azide at 4 °C. Sections were incubated in 5% horse serum in 0.5% Triton-X TBS for 1 hour before overnight incubation with 1:500 anti-GFP or 48 hours 1:300 anti-UBE3A (3E5 Sigma) and 1 hr 1:500 Alexa-Fluor 488 (Thermo Fisher Scientific, A-11039) or Cy3-conjugated (Jackson Immunoresearch, 715-165-151) secondary, respectively. Sections were stained with Hoechst for nuclei labeling before mounting in Fluoromount-G (Southern Biotech).

### Ex vivo dendritic spine analysis

Images in a z series projection of three areas of each neuron (basal, proximal and distal) was taken at 63x. To measure spine density, NeuronStudio (Rodriguez et al., 2008) was used to automatically define neurites and quantify spine number and subtype (thin, mushroom, or stubby) for each neuron image. Per the automated software, thin and mushroom spines were first defined as having a head to neck ratio greater than 1.1. To be further classified as a thin spine if this neck ratio was not met, spines were required to have a length to spine head ratio of 3. Mushroom spines were further defined as having a head diameter of greater than 0.3µm in addition to the neck ratio of 1.1 or more. Otherwise, spines were classified as stubby. Density was calculated as spines per micron per region per neuron, with multiple neurons per animal. Total (F (3, 32) = 21.38). Oriens (F (3, 32) = 33.59). Radiatum (F (3, 32) = 38.57). Moleculare (F (3, 32) = 21.81).

### Electrophysiology recordings

P30 129S7 mice were anesthetized with Euthasol and decapitated. Brains were quickly removed and placed in ice-cold ACSF. Sagittal sections (300 µm) containing hippocampus were prepared in ice-cold ACSF using a vibrating blade microtome (Leica VT1200). Right after cutting, slices were recovered for 10 minutes at 32 °C and then transferred to holding ACSF at room temperature. Cutting and recovery were performed with ACSF containing the sodium substitute NMDG (Ting et al., 2014): 92 mM NMDG, 20 mM HEPES (pH 7.35), 25 mM glucose, 30 mM sodium bicarbonate, 1.2 mM sodium phosphate, 2.5 mM potassium chloride, 5 mM sodium ascorbate, 3 mM sodium pyruvate, 2 mM thiourea, 10 mM magnesium, 14 mM sulfate, 0.5 mM calcium chloride. ACSF used for holding slices prior to recording was identical but contained 92 mM sodium chloride instead of NMDG and 1 mM magnesium chloride and 2 mM calcium chloride. ACSF used to perfuse slices during recording contained: 125 mM sodium chloride, 2.5 mM potassium chloride, 1.25 mM sodium phosphate, 1 mM magnesium chloride, 2.4 mM calcium chloride, 26 mM sodium bicarbonate, and 11 mM glucose. All ACSF solutions were saturated with 95% O2 and 5% CO2. For recording, a single slice was transferred to a heated chamber (32 °C) and perfused with normal ACSF (2.5 ml min−1) using a peristaltic pump (WPI). Visualization of neurons in the pyramidal layer of hippocampal CA1 was performed with an upright microscope equipped for differential interference contrast (DIC) microscopy (BX51WI, Olympus). Whole-cell patch-clamp recordings were made using a MultiClamp 700B amplifier (1 kHz low-pass Bessel filter and 10 kHz digitization) with pClamp 10.3 software (Molecular Devices). Voltage-clamp recordings were made using glass pipets with resistance 3-5 MOhms, filled with internal solution containing: 117 mM cesium methanesulfonate, 20 mM HEPES, 0.4 mM EGTA, 2.8 mM NaCl, 5 TEA-Cl, 2.5 mM Mg-ATP, and 0.25 mM Na-GTP, pH 7.2–7.3 and 285 mOsm. Input resistance was continually monitored on-line; cells with input resistance changes greater than 20% were not included in the analysis. mEPSCs were pharmacologically isolated by having tetrodotoxin (1 μM), gabazine (1 μM), and APV (50 μM) present throughout the experiment and sampled at 1.8 KHz while clamping the cells at −70 mV. 200-300 events per cell were analyzed, using a threshold of 2X the baseline noise. Analysis of mEPSCs was performed off-line using the MiniAnalysis program (v 6.0). For evoked EPSC experiments, electrical stimuli were delivered in the presence of gabazine (1 μM) to the Schaffer collaterals (stratum radiatum of CA1) at 0.1 Hz via a bipolar electrode. Paired pulse ratios were acquired at −70 mV, by having a second afferent stimulus of equal intensity at several intervals (20-500 ms) after the initial stimulus. The ratio was calculated by dividing the peak amplitude of the second stimulus with the peak amplitude of the first stimulus. Input output curves were obtained by delivering stimuli at a range of intensities (20-500 μAmp) to each cell and measuring peak amplitudes of evoked EPSCs. For mEPSC frequency and amplitude and rise and decay times, number of cells 129S7: WT (n=10), AS (n=13), AS/E5^-/-^ (n=12), E5^-/-^ (n=9). Frequency (F(3, 40) = 6.779). Amplitude (F(3, 40) = 2.757). Rise time (F (3, 40) = 6.127. Decay time F (3, 40) = 3.337. For I/O and PPF, number of cells 129S7: WT (n=14), AS (n=12). I/O Interaction (F (9, 238) = 2.731), Stimulus Intensity (F (9,238) = 36.46). Genotype (F (1, 238) = 29.93). PPF Interaction (F (4, 118) = 0.9733), Interstimulus Interval (F (4, 118) = 44.34), Genotype (F (1, 118) = 0.6969). 129S7: I/O: WT (n=11), AS/E5^-/-^ (n=12). Interaction (F (9, 210) = 1.753), Stimulus Intensity (F (9, 210) = 22.10). Genotype (F (1, 210) = 22.34). PPF: WT n=10), AS/E5^-/-^ (n=12). Interaction (F (4, 100) = 0.3177), Interstimulus Interval (F (4, 100) = 27.36), Genotype (F (1, 100) = 2.535).

### Transfection

At DIV10, neurons were transfected with plasmids containing GFP to visualize dendritic processes and spines, shRNA hairpin to UBE3A or a scrambled version to knockdown the indicated protein, or HM4D(G_i_), to modulate activity, using Lipofectamine 2000. In short, the media on the neurons was reduced and media containing a mixture of Lipofectamine 2000 and the plasmid DNA was added for 1 hour. The media was completely removed and replaced with the previously removed media.

### DREADD drug treatment

At DIV11, neurons were treated with a final concentration of 10 μM CNO dissolved in DMSO or equivalent amounts of DMSO directly into the media. This concentration was maintained through feedings until coverslips were taken for fixing and staining.

### Immunocytochemistry

Coverslips were paraformaldehyde fixed in PBS before 48 hour incubation in 1:300 mouse anti-UBE3A and 1:300 chicken anti-GFP followed by 1 hr 1:500 Cy3-conjugated anti-mouse secondary and 1:500 488-conjugated anti-chicken secondary. Neurons were incubated with Hoechst for nuclei labeling before mounting in Fluoromount-G.

### In vitro spine imaging

Images in a z series projection of single neurons were taken at 40x and averaged over 2 images. To measure spine density, NeuronStudio (Rodriguez et al., 2008) was used to automatically define neurites and quantify spine number for each neuron image. Density was calculated as spines per micron.

### Quantification and statistical analysis

Statistical analysis was performed using Graphpad Prism 6. All tests were two-tailed and *p* values ≤ 0.05 were considered significant. N for each experiment, degrees of freedom, and f/t values are listed in figures or materials and methods. Statistical tests and *p* values for indicated figures and post-hoc tests and *p* values for indicated figures are shown in **Table 2** and **Table 3**.

**Table 2:** Statistical tests and p values for indicated Figures.

| Figure | Statistical Test | P value | Significance |
| --- | --- | --- | --- |
| 1B – Ephexin5 levels<br>P91 | Student's T-test | 0.0094 | * |
| 1C - NPP | Two-way ANOVA | Object: p=0.0046<br>Genotype: p > 0.9999<br>Interaction: p < 0.0001 | *<br>ns<br>* |
| 1D - NOR | Two-way ANOVA | Object: p<0.0001<br>Genotype: p>0.9999<br>Interaction: p=0.0620 | *<br>ns<br>ns |
| 1E - Rotarod | One-way ANOVA | p < 0.0001 | * |
| 1F - Rotarod | One-way ANOVA | p=0.0006 | * |
| 1G – Marble burying | One-way ANOVA | p=0.0002 | * |
| 2B – Ephexin5 and<br>UBE3A levels in Cre<br>animals | Student's T-test, Welch's<br>correction | Ephexin5: 0.0016<br>UBE3A: 0.0473 | *<br>* |
| 2C – Rotarod | Student's T-test | 0.5509 | ns |
| 2D - NPP | One-way ANOVA<br>Kruskal-Wallis<br>One-way ANOVA<br>Kruskal-Wallis | p=0.0002<br>p=0.2867<br>p=0.0004<br>p=0.0039 | *<br>ns<br>*<br>* |
| 3C – Spine density | One-way ANOVA | p < 0.0001 | * |
| 4B – mEPSC<br>frequency | One-way ANOVA | p=0.0008 | * |
| 4C – mEPSC<br>amplitude | One-way ANOVA | p=0.0548 | ns |
| 4E – Rise time | One-way ANOVA | p=0.0016 | * |
| 4F – Decay time | One-way ANOVA | p=0.0287 | * |
| 1-1D – Ephexin5 levels<br>P30 | Student's T-test | p=0.0722 | ns |
| 1-1E – Passive<br>avoidance | Two-way ANOVA | Genotype: p=0.0016<br>Day: p < 0.0001<br>Interaction: p=0.0012 | *<br>*<br>* |
| 1-1F – Passive<br>avoidance | Two-way ANOVA | Genotype: p=0.6671<br>Day: p<0.0001<br>Interaction: p=0.6671 | ns<br>*<br>ns |
| 3-1B – Spine density | One-way ANOVA | p < 0.0001 | * |
| 3-1E – Spine density | One-way ANOVA | p < 0.0001 | * |
| 3-1H – Spine density | One-way ANOVA | p < 0.0001 | * |
| 4-1C – Input/Output | Two-way ANOVA | Stimulus: p < 0.0001<br>Genotype: p < 0.0001 | *<br>* |
|  |  | Interaction: 0.0177 | * |
| 4-1D - PPF | Two-way ANOVA | ISI: p<0.0001<br>Genotype:0.4055<br>Interaction: p=0.4170 | *<br>ns<br>ns |
| 4-1F – Input/Output | Two-way ANOVA | Stimulus:p<0.0001<br>Genotype:p<0.0001<br>Interaction: p=0.0789 | *<br>*<br>ns |
| 4-1H - PPF | Two-way ANOVA | ISI: p<0.0001<br>Genotype: 0.1145<br>Interaction: 0.8656 | *<br>ns<br>ns |
| 5B – Spine density WT | Two-way ANOVA | Interaction: p=0.0886<br>Hairpin: p=0.4370<br>Drug: p=0.0260 | ns<br>ns<br>* |
| E5 | Two-way ANOVA | Interaction: p=0.9530<br>Hairpin: p=0.0685<br>Drug: p=0.4835 | ns<br>ns<br>ns |

**Table 3:**
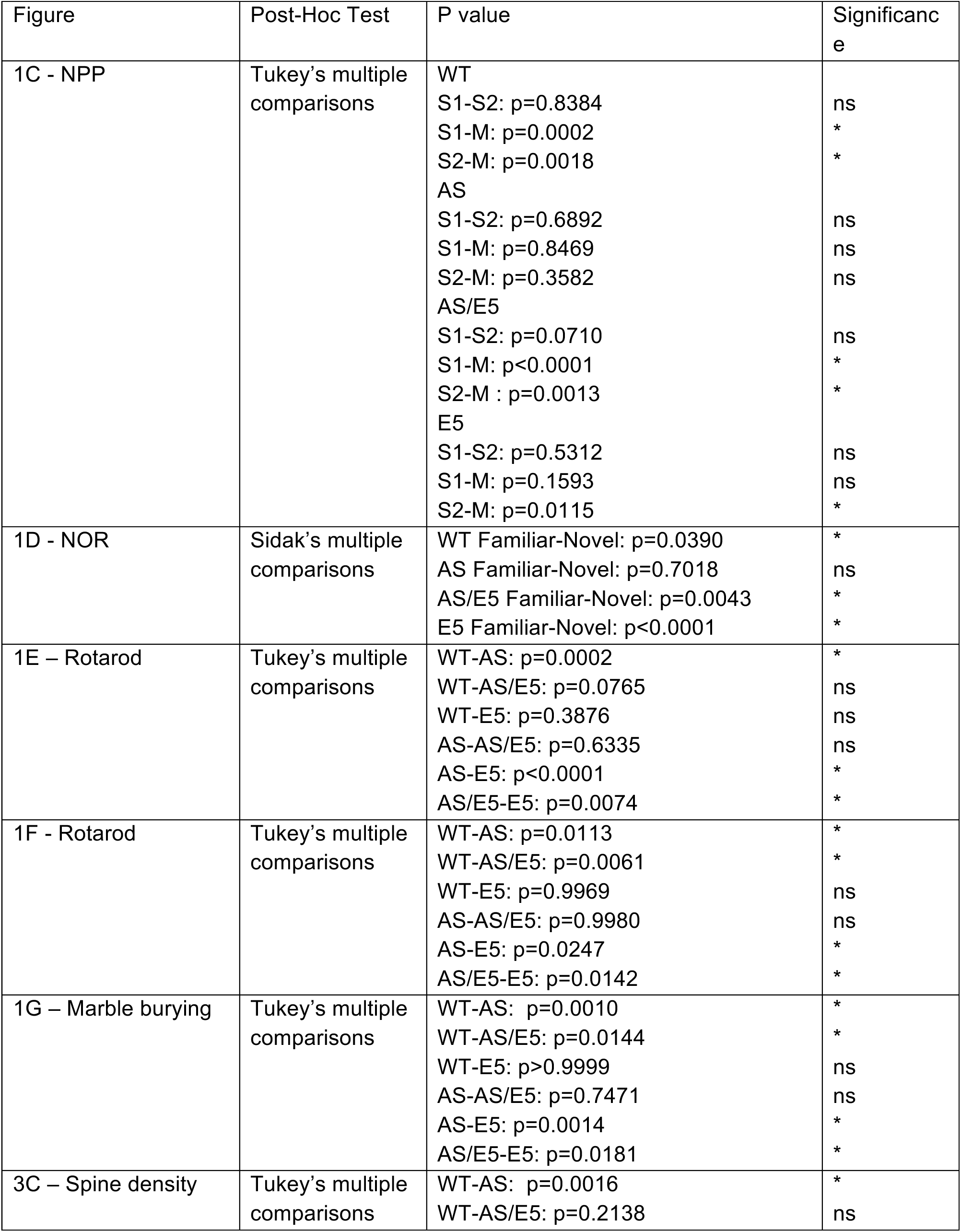
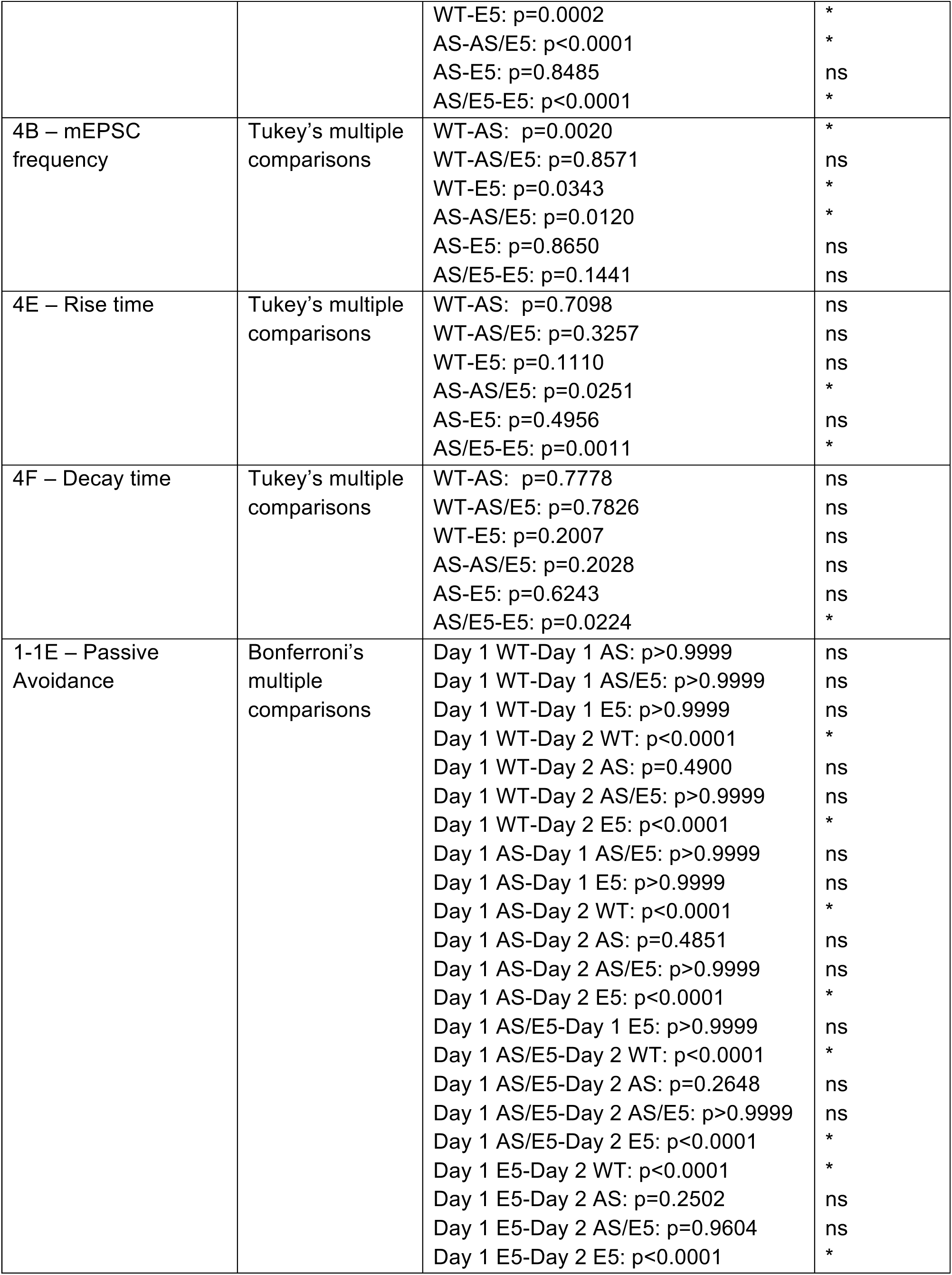
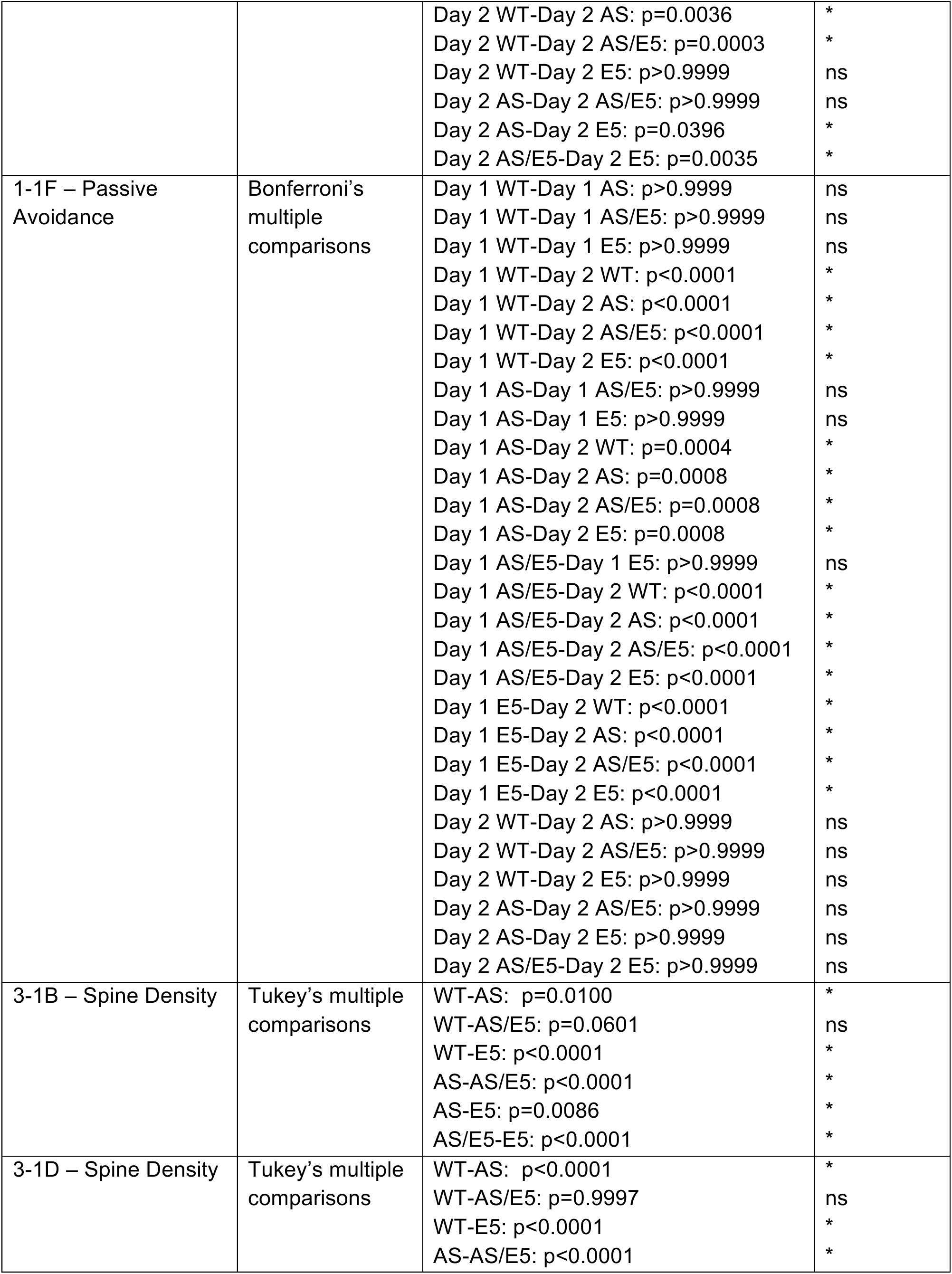
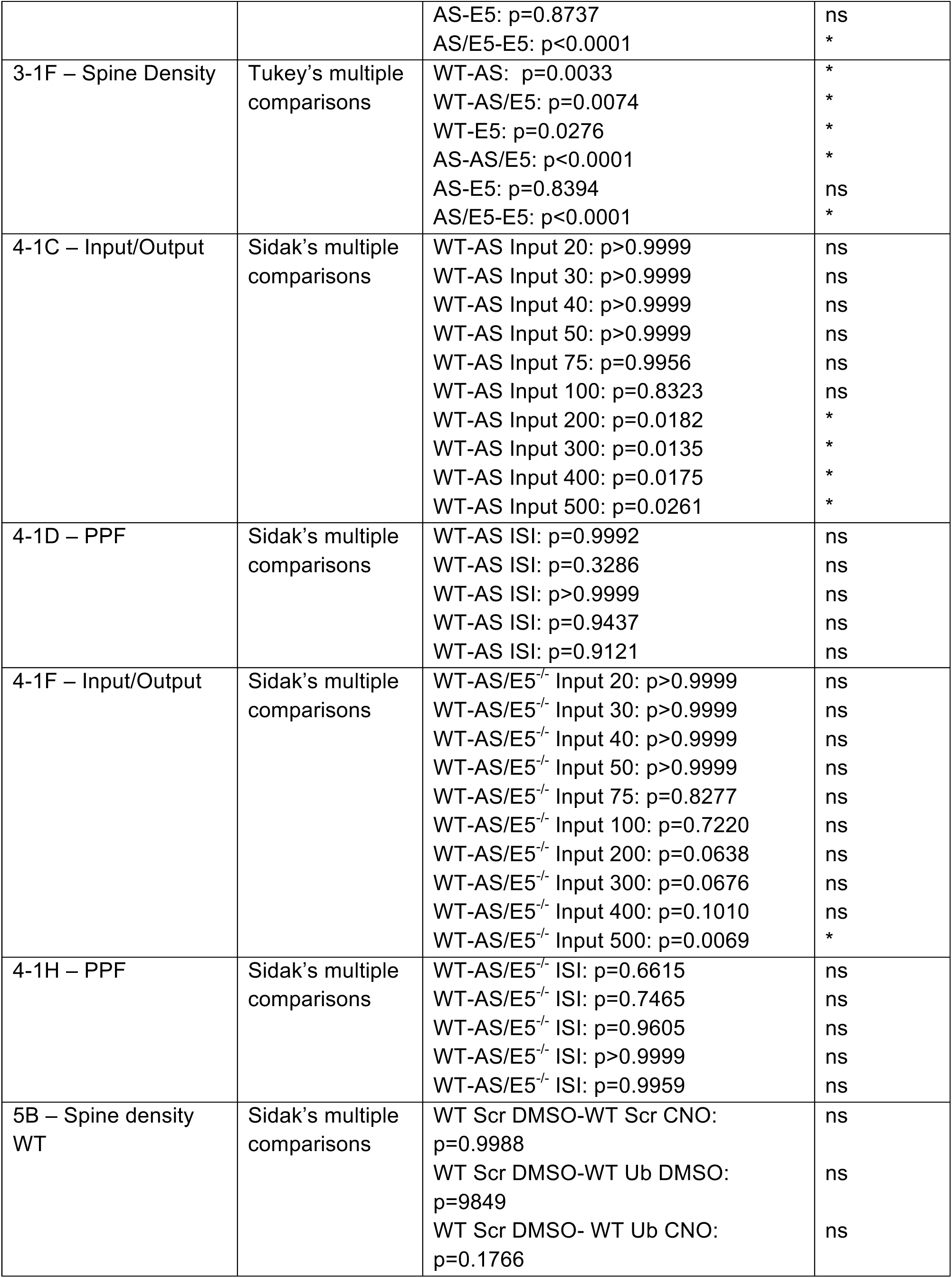
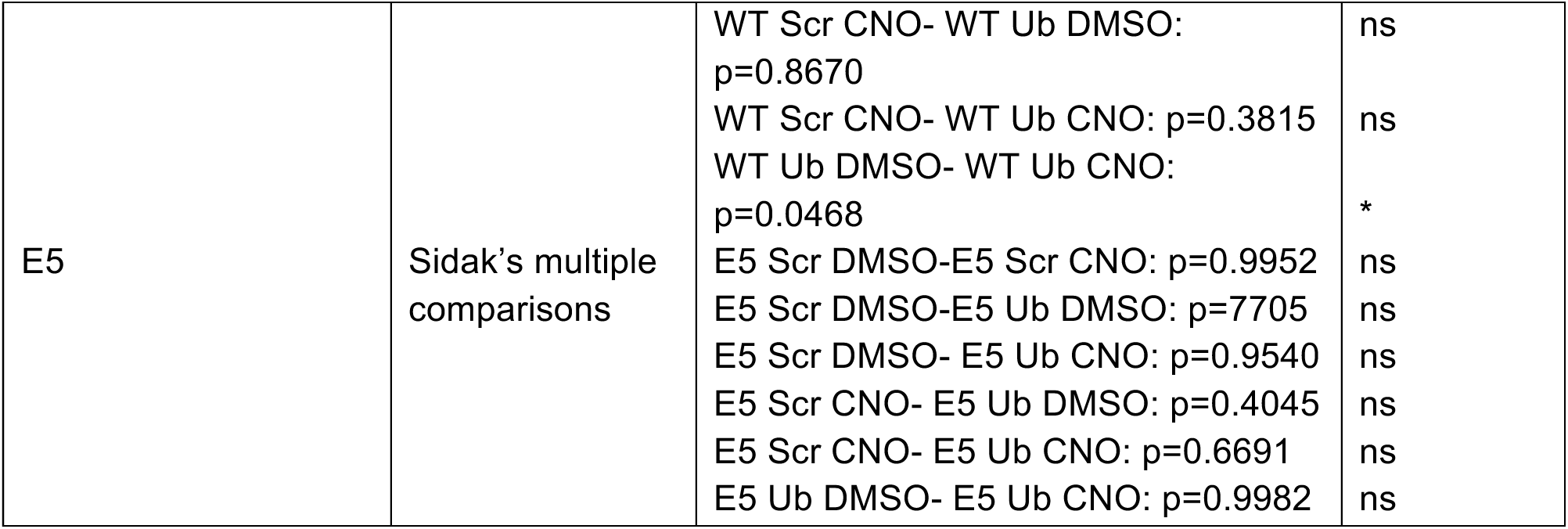
Post-hoc tests and p values for indicated figures.

## RESULTS

### Ephexin5 is a substrate of UBE3A *in vitro* and Ephexin5 expression is increased *in vivo* in the hippocampus of AS mouse models

Previous reports have demonstrated that Ephexin5 is ubiquitylated in brains of mice, which is, in part, dependent on the presence of UBE3A (Margolis et al., 2010). To determine whether Ephexin5 can be directly ubiquitylated by UBE3A, we used purified Ephexin5 protein and incubated it in an *in vitro* purified UBE3A ubiquitin ligase assay. Samples were then run on SDS-PAGE for immunoblot analysis using antibodies raised against Ephexin5. We noted a shift in Ephexin5 mobility consistent with ubiquitin modification (Fig. 1*A*), which was dependent upon the presence of ATP and UBE3A (Extended Data Fig.1-1*A*). E6 protein was included in the initial experiment to get enhanced ubiquitylation of Ephexin5 since it promotes UBE3A activity (Ronchi et al., 2014). While E6 protein enhances this in vitro assay, it is not required for UBE3A to ubiquitylate Ephexin5 at several residues (Extended Data Fig.1-1*A*). We confirmed that Ephexin5 was directly ubiquitylated by mass spectrometry analysis and identified UBE3A catalyzed ubiquitylation sites on Ephexin5 with and without E6 (Extended Data Fig.1-1*B,C*). Taken together, these data identify Ephexin5 as a highly likely substrate of UBE3A-dependent ubiquitylation. Consistent with the idea that this ubiquitylation targets Ephexin5 for proteasome degradation, previous findings have shown that Ephexin5 expression is elevated early in postnatal AS mouse brain (Margolis et al., 2010). To determine whether Ephexin5 levels were also elevated in adult hippocampus, when hippocampal-dependent behaviors and hippocampal synaptic transmission are known to occur, we measured hippocampal Ephexin5 levels in wild type (WT) and AS littermates of the 129S7 strain at postnatal day 91 (P91). Dissected hippocampi were lysed and prepared for SDS-PAGE and western analysis using antibodies raised against UBE3A, Ephexin5, or Actin. We detected a significant elevation in Ephexin5 antibody signal in AS mice as compared to WT littermate controls at this age (Fig. 1*B*).

**Figure. 1.**
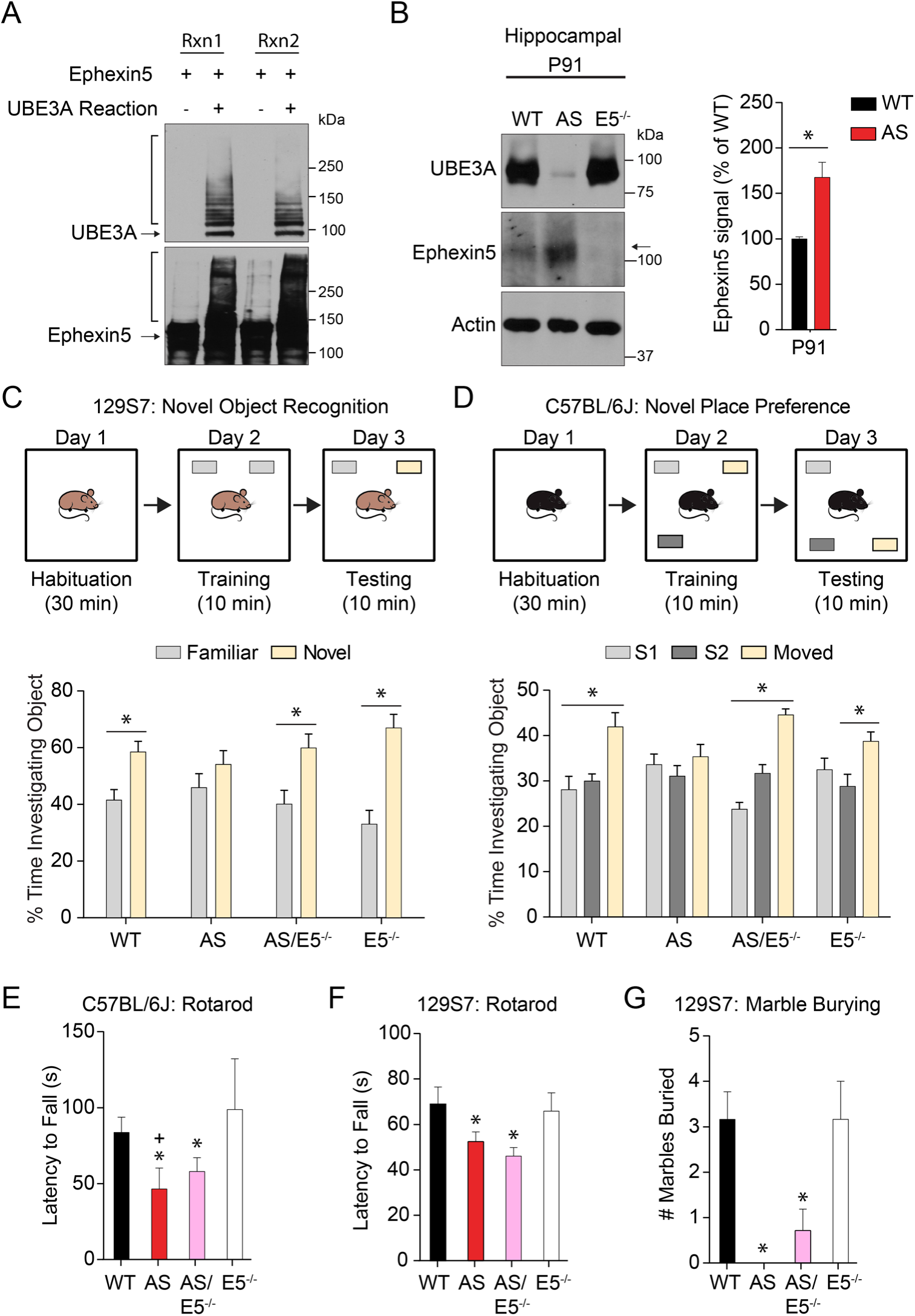
Ephexin5 is upregulated in AS mouse hippocampal tissue and removal of Ephexin5 modulates learning and memory behavioral task. **(A)** *In vitro* ubiquitylation assays were performed using purified Ephexin5 protein in the presence or absence of UBE3A reaction buffer (containing UBE1, UBE2L3, UBE3A, E6, Ubiquitin, and Mg-ATP), as indicated. Replicates shown using immunoblot analysis with indicated antibodies. The arrows indicate the unmodified protein and the bracket indicates the smearing due to ubiquitylation. **(B)** Ephexin5 expression levels are elevated in AS mouse hippocampus. Hippocampi were dissected from wild-type (WT) and AS mice at P91 and lysates were prepared for immunoblotting with indicated antibodies. Quantification of Ephexin5 signal is normalized to Actin signal and compared to WT. Arrow indicates Ephexin5 band. Data are presented as mean ± SEM (n = 4 for WT, n = 5 for AS). \**p*<0.05 (unpaired Student’s t-test). **(C)** Object investigation in the NPP test in 3 month-old C57Bl/6j mice. Shown is the percent time spent investigating each object. Data are presented as mean ± SEM. \**p*<0.05 (two-way ANOVA) compared to the stationary objects (S1 and S2) or as indicated with *post-hoc* Bonferroni multiple comparisons test. **(D)** Object investigation in the NOR test in 3 month old 129S7 mice. Shown is the percent time spent investigating each object. Data are presented as mean ± SEM. \**p*<0.05 (two-way ANOVA) compared to familiar within genotype with *post-hoc* Holm-Sidak correction. **(E)** Latency for C57Bl/6j mice to fall off the rotating rod, measured in seconds. Data are presented as mean ± SEM. \**p*<0.05 compared to E5^-/-^, ^+^*p*<0.05 compared to WT (one-way ANOVA) with *post-hoc* Tukey’s multiple comparisons test. **(F)** Latency for 129S7 mice to fall off the rotating rod, measured in seconds. Data are presented as mean ± SEM. \**p*<0.05 (one-way ANOVA) compared to WT and E5^-/-^with *post-hoc* Tukey’s multiple comparisons test. **(G)** Number of marbles buried or covered 2/3 in 20 minutes by 129S7 mice. Data are presented as mean ± SEM. \**p*<0.05 compared to WT and E5^-/-^ (one-way ANOVA) with *post-hoc* Tukey’s multiple comparisons test. See also **Extended Data Fig.1-1**. Sample size (n), degrees of freedom, and exact *p* values are reported in **Table 2** and **Table 3**.

### Removal of Ephexin5 in the AS mouse model rescues hippocampus-dependent behaviors

To determine whether elevated levels of Ephexin5 play a crucial role in AS-relevant phenotypes, we used genetic approaches to delete Ephexin5 in the 129S7 AS mouse model. Breeding strategies were devised to generate wild type (WT), Angelman Syndrome (*Ube3A^m-/p^*^+^, AS), Ephexin5 knockout (*Ephexin5^-/-^*, E5^-/-^) and double knockout (*Ube3A^m-/p+^/Ephexin5^-/-^*, AS/E5^-/-^). Ephexin5’s role in the nervous system has been predominately described in the hippocampus where its expression is enriched (Margolis et al., 2010). Therefore, we tested WT, AS, AS/E5^-/-^, and E5^-/-^ littermates on several hippocampus-dependent and hippocampus-independent behavioral tasks (Barker and Warburton, 2011). We used a traditional novel object recognition task (NOR) and passive avoidance to investigate hippocampus-dependent behaviors. While the AS mouse showed a lack of preference for the novel object (Fig. 1*C*), there was no difference in the response of any genotype within the passive avoidance (Extended Fig. 1-1*D*). However, these passive avoidance results are difficult to interpret, as none of the animals entered the dark arena on test day within the 5-minute test trial. Importantly, removal of Ephexin5 in the AS mouse model was sufficient to restore preference for the novel object (Fig. 1*C*).

Previous studies indicate UBE3A deletion on pure 129S7 vs C57Bl/6J have different phenotype penetrance (Born et al., 2017; Huang et al., 2013b; Sittig et al., 2016). We sought to extend our observation in 129S7 strain into C57BL/6J to determine the general relevance of the Ephexin5 pathway in AS-associated hippocampal dependent phenotypes. We generated WT, AS, AS/E5^-/-^, and E5^-/-^ littermates on C57Bl/6J. At P30, we noted an increase in Ephexin5 levels in the AS mice (Extended Data Fig. 1-1*E*). We chose to test these mice in the novel place preference test (NPP) which our lab has extensive experience with (Sell et al., 2017) and is a similar task to NOR known to rely on the same brain area (Stackman et al., 2016). The AS animals did not prefer the moved object as measured by the amount of time spent investigating each object 24 hours later, in contrast to the WT, AS/E5^-/-^, and E5^-/-^ animals which showed a significant preference for the moved object (Fig. 1*D*). Interestingly, in the passive avoidance task, which requires both hippocampus and amygdala activity (van der Poel, 1967), Ephexin5 deletion did not rescue C57Bl/6J AS mouse behavior (Extended Data Fig.1-1*F*).

In contrast to these more hippocampal specific learning tasks, Ephexin5 deletion did not improve AS mouse performance in the rotarod motor coordination task in either mouse strain (Fig. 1*E-F*) and the marble-burying task in the 129S7 strain (Huang et al., 2013b) (Fig. 1*G*). This lack of rescue may be due to the regionally restricted expression and activity of Ephexin5, as neither the rotarod nor the marble-burying task is dependent on hippocampal function. These data provide evidence for regulation of hippocampal function by UBE3A in behaviorally relevant ways, which is at least in part dependent on the presence of Ephexin5. Moreover, we concluded that removal of Ephexin5 does not interfere with hippocampus-dependent learning or other behaviors in a wild-type background. Thus, removal of Ephexin5 appears to specifically rescue hippocampus-dependent learning deficits observed in AS mice, indicating that the elevation of Ephexin5 observed in AS mice may be pathological.

### Conditional removal of UBE3A in excitatory forebrain neurons recapitulates learning and memory but not motor phenotypes of AS mice

UBE3A is expressed throughout the nervous system in a variety of cell types, whereas Ephexin5 is most highly expressed in excitatory neurons within the hippocampus (Margolis et al., 2010). To determine whether UBE3A within hippocampal excitatory cells was responsible for the cognitive behavioral deficits observed in AS mice, we used the Cre/Lox system to conditionally remove UBE3A from the C57Bl/6J:ube3A^loxp^ (Judson et al., 2016). In order to obtain robust, consistent recombination in CA1 and DG, we used the CamKIIa-Cre line, which is expressed in excitatory forebrain neurons by 2 weeks postnatally and produces robust recombination in hippocampus (McGill et al., 2018). To identify the cells in which UBE3A expression was removed, we crossed the CamKIIa-Cre line with a TdTomato reporter line. At 8 weeks, expression of the reporter was detected in a subset of excitatory cortical neurons and robustly in the excitatory cell layers of CA1 and DG (Fig, 2*A*). Lysates from hippocampi of adult Cre^+^ and Cre^-^maternal Ube3A^loxp+^ were analyzed via Western blot for expression of UBE3A and Ephexin5. Compared to the hippocampi of Cre^-^animals, in the hippocampi of Cre^+^ animals UBE3A expression was significantly decreased, while Ephexin5 was significantly increased (Fig. 2*B* and Extended Data Fig. 2-1). Consistent with previous descriptions of Ephexin5 expression, in Cre^-^animals expression of Ephexin5 in the cortex was undetectable as compared to hippocampus (Extended Data Fig. 2-1). Nevertheless, to determine whether the CamKIIa-Cre expression in a subset of cortical neurons produces deficits in cortex-dependent behaviors in UBE3a-Floxed mice, we tested animals on the rotarod task, which requires cortical function (Bruinsma et al., 2015). Expression of Cre in this subset of cortical neurons did not decrease the latency to fall from the rotating rod (Fig. 2*C*), as was measured in AS animals (Fig. 1*E*). In contrast, deficits in the novel place preference task were observed in Cre^+^ animals (Fig. 2*D*), similar to the AS animals (Fig. 1*C*). Furthermore, this deficit was again rescued by removal of Ephexin5 (Fig. 2*D*), suggesting that Ephexin5 most likely acts within the hippocampus to ameliorate these deficits.

**Figure. 2.**
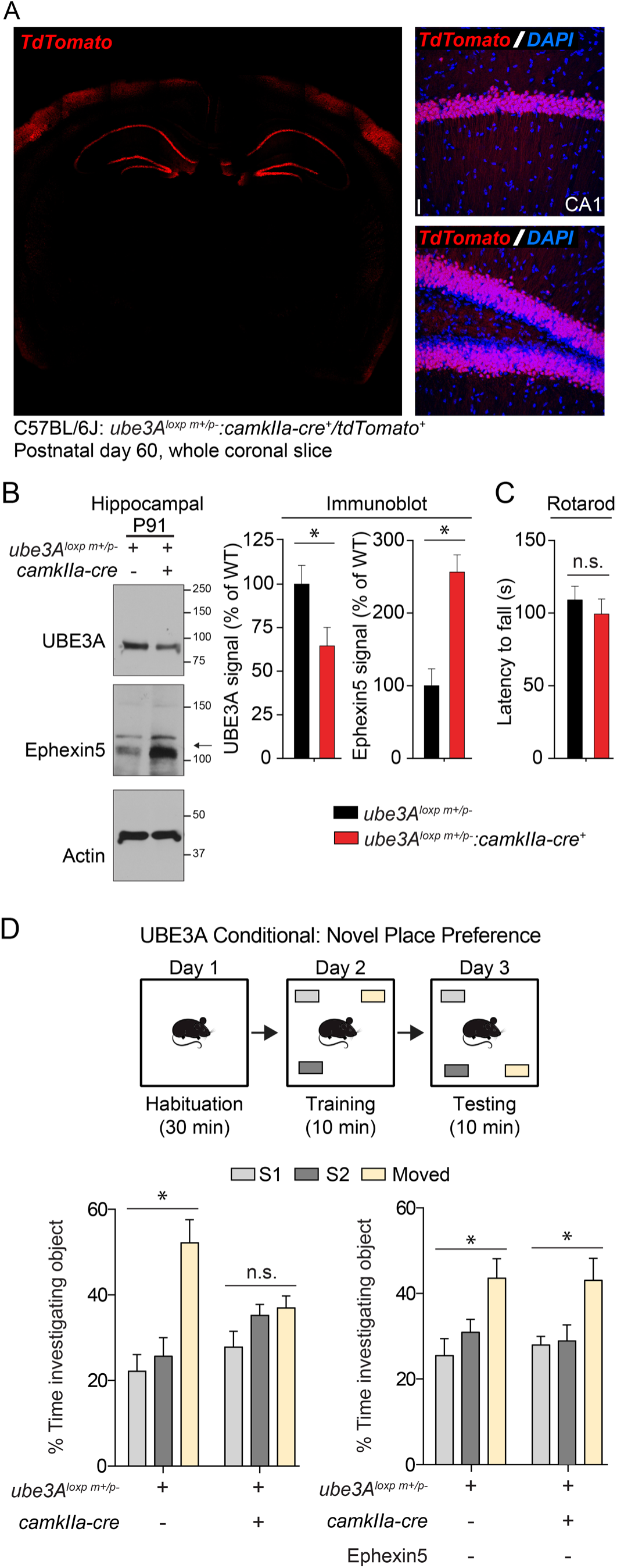
*Ube3A^loxp+^/camkIIa-cre^+^* mice have increased Ephexin5 expression in the hippocampus and Ephexin5 dependent deficits in NPP. **(A)** 8 week old TdTomato cre reporter-positive, *camkIIa-cre*^+^ tissue was stained for TdTomato (red) with Hoechst (DAPI - blue) for nuclei labeling. On the left is a 10x image of one coronal section indicating strong hippocampal and some cortical expression. On the right are images of DG and CA1, indicating nearly 100% recombination in the excitatory cell layers. Scale bar = 10 µm **(B)** Ephexin5 expression levels are elevated in hippocampus from *ube3A^loxp+^/camkIIa-cre^+^* mice. Quantification of Ephexin5 and UBE3A signal is normalized to Actin signal and compared to *ube3Aloxp^+^/camkIIa-cre^-^* controls. Arrow indicates Ephexin5 band. Data are presented as mean ± SEM (n = 6 for *ube3A^loxp+^/camkIIa-cre^+^*, n = 5 for *ube3A^loxp+^/camkIIa-cre*^-^). * *p*<0.05 (unpaired Student’s t-test). **(C)** Latency for WT C57Bl/6j mice, with or without *camkIIa-cre* dependent *ube3A* removal, to fall off the rotating rod, measured in seconds. **(D)** Object investigation in the NPP test in 3 month-old C57Bl/6j mice, with or without *camkIIa-cre* dependent *ube3A* removal in a WT or E5^-/-^ background. Shown is the percent time spent investigating each object. Data are presented as mean ± SEM. \**p*<0.05 (two-way ANOVA) compared to the stationary objects (S1 and S2) with *post-hoc* Bonferroni multiple comparisons test. See also **Extended Data Fig. 2-1**. Sample size (n), degrees of freedom, and exact *p* values are reported in **Table 2** and **Table 3**.

### Removal of Ephexin5 restores hippocampal CA1 spine density in AS mice to WT levels

Previous studies in visual cortex and striatum have reported deficits in synaptic transmission in AS mice (Hayrapetyan et al., 2014; Kim et al., 2016; Wallace et al., 2012), as well as decreases in spine density in AS mice using Golgi staining or phalloidin staining in CA1 (Dindot et al., 2008; Kim et al., 2016). We suspected that, since Ephexin5 is involved in developmental regulation of dendritic spine formation, AS mice may also have altered spine morphology in the hippocampus. Furthermore, novel object recognition and object location recall have been linked specifically to hippocampal CA1 function (Haettig et al., 2013; Stackman et al., 2016). To test this idea, we included the Thy1-EGFP (enhanced green fluorescent protein expressed from the Thy1 promoter) line in our crosses that generate WT, AS, AS/E5^-/-^ and E5^-/-^ lines to label a subset of neurons in the hippocampus with EGFP. At 11 weeks of age, the Thy1-EGFP labeled lines were anesthetized and perfused. Brains were then sectioned and imaged to visualize labeled pyramidal neurons. We focused on the CA1 region of hippocampus, where UBE3A is highly expressed (Extended Data Fig. Fig. 3-1*A*). We visualized dendritic spines on individual EGFP-expressing neurons using confocal microscopy (Extended Data Fig. Fig. 3-1*B*). Dendrites from each CA1 pyramidal neuron were imaged and spine densities quantified from the stratum radiatum (s.r.), stratum oriens (s.o.) and stratum lacunosum-moleculare (s.l.m) (Fig. 3 A,B,E,H). Changes in dendritic spine density were consistent across strata of imaged CA1 pyramidal neurons and thus combined in the analysis. Unexpectedly, blinded analysis revealed an increase in spine density on CA1 pyramidal neurons of AS mice. This is in contrast to a previous study which reported, using Golgi staining, decreased dendritic spine density in the CA1 of the AS hippocampus (Dindot et al., 2008). In our study, the relative abundance of spine types (stubby, thin, mushroom) did not differ significantly between AS and WT mice (Fig. 3C,D,F,G,I,J). Consistent with previous ex *vivo* studies in developing E5^-/-^ animals (Hamilton et al., 2017; Margolis et al., 2010), we observed *ex vivo* that dendrites from adult E5^-/-^ animals also showed an increase in spine density compared to WT controls. Importantly, deletion of Ephexin5 in the AS/E5^-/-^ mice restored dendritic spine density to WT levels. Looking at the distribution of spine type, as measured by length and width of each dendritic spine counted, we observed that the proportions of spine subtypes from the AS/E5^-/-^ mice were altered compared to other genotypes, with an increase in stubby spines and a decrease in thin spines (Fig. 3 and Extended Data Fig. 3). This change may be attributed to a consistent decrease in AS/E5^-/-^ thin spines compared to AS and E5 animals. Stubby spines are most common in early postnatal development, after a loss of filopodia and before a large increase in thin and mushroom spines (Harris et al., 1992). This decrease in thin spines may indicate that a larger subset of the spines in the AS/E5^-/-^ animals are less mature or are developing later than spines in the other genotypes (Hering and Sheng, 2001). Taken together, these data identify a unique feature of AS mice, whereby in the CA1 there is an increase in dendritic spine density across strata compared to WT mice that is at least in part dependent on the presence of Ephexin5.

**Figure. 3.**
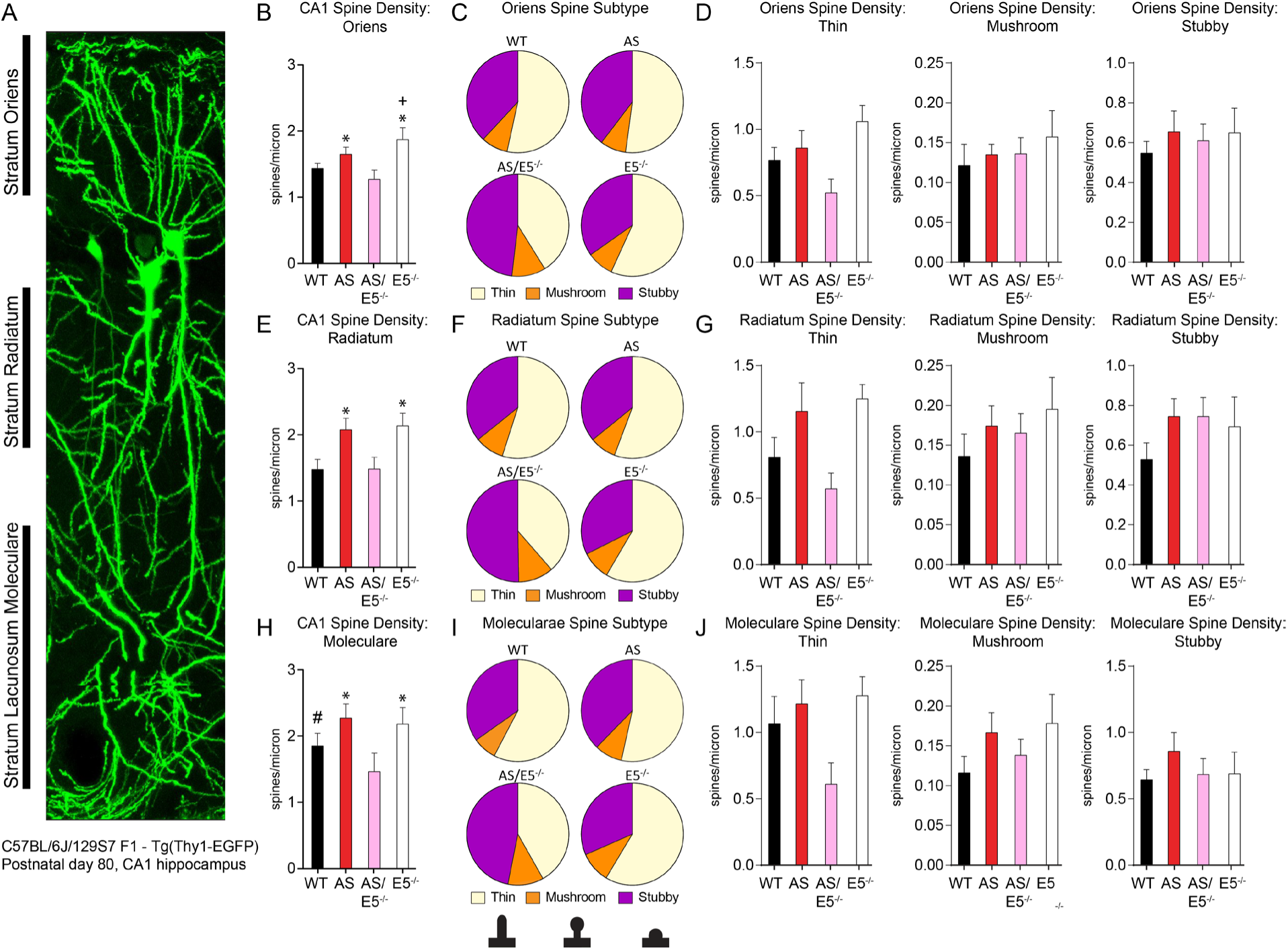
UBE3A expression in CA1 and location of spine analysis in CA1 GFP-positive neurons, related to Figure 3. **(A)** Example CA1 GFP-positive neurons with the regions of interest labeled (stratum oriens, stratum radiatum, and stratum moleculare). Scale bar is 10 μm. **(B)** Oriens spine density in CA1. **(C)** Proportion of spine subtypes within WT, AS, AS/E5^-/-^, and E5^-/-^ oriens dendrites of CA1 neurons. **(D)** Spine density in the stratum oriens of CA1 neurons for thin, mushroom, and stubby morphologies. **(E)** Radiatum spine density in CA1. **(F)** Proportion of spine subtypes within WT, AS, AS/E5^-/-^, and E5^-/-^ radiatum dendrites of CA1 neurons. **(G)** Spine density in the stratum radiatum of CA1 neurons for thin, mushroom, and stubby morphologies. **(H)** Moleculare spine density in CA1. **(I)** Proportion of spine subtypes within WT, AS, AS/E5^-/-^, and E5^-/-^ moleculare dendrites of CA1 neurons. **(J)** Spine density in the stratum lacunosum moleculare of CA1 neurons for thin, mushroom, and stubby morphologies. One-way ANOVA, *post-hoc* Tukey’s multiple comparisons test for panels B, E, and H. Two-way ANOVA, *post-hoc* Sidak’s multiple comparisons test for panels D, G, and J. All data are presented as mean ± SEM (n = 3 for all genotypes) \**p*<0.05 compared to WT and AS/E5^-/-^, ^+^*p*<0.05 compared to AS, ^#^*p*<0.05 compared to AS/E5^-/-^ for all panels. See also **Extended Data Fig. 3-1**. Degrees of freedom and exact *p* values are reported in **Table 2** and **Table 3**.

### Removal of Ephexin5 restores CA1 excitatory transmission in AS mice to WT levels

To determine the physiological consequence of the observed changes in dendritic spine density described in Fig. 3, we recorded miniature excitatory postsynaptic currents (mEPSCs) from pyramidal neurons in the CA1 (Fig. 4*A*). To measure number and strength of excitatory synapses in individual neurons, mEPSCs were isolated at −70mV in the presence of tetrodotoxin (TTX, 1 μM), gabazine (1 μM), and APV (30 μM). Consistent with the observed increase in spine density, AS mice exhibited a higher frequency of mEPSCs than WT littermates (Fig. 4*B*); mEPSC amplitudes were not statistically different (Fig. 4*C*). Stimulus-evoked EPSCs were also significantly larger in AS mice (Extended Data Fig. 4-1*A-B*). This increase in excitatory inputs does not appear to result from a change in presynaptic glutamate release, as paired pulse ratios – a measure of presynaptic release probability – across a range of pulse intervals were not different between AS and WT mice (Extended Data Fig. 4-1*C-D*). Consistent with previous reports (Margolis et al., 2010), E5^-/-^ animals also exhibited higher mEPSC frequency relative to WT (Fig. 4*B*). However, mEPSC frequency did not differ between WT and AS/E5^-/-^ mice (Fig. 4*B*). The average rise time and decay time of miniature events were slightly longer in AS/E5^-/-^ mice, though not statistically different from WT (Fig. 4*E-F*). To determine whether evoked excitatory inputs were also similar between AS/E5^-/-^ animals and WT animals, stimulus-evoked EPSCs were recorded from the CA1 of AS/E5^-/-^ and WT littermate controls (Extended Data Fig. 4-1*E-F*). In contrast to AS animals, AS/E5^-/-^ animals exhibited only a slight decrease in EPSC amplitude at the highest stimulus intensity (Extended Data Fig. 4*A-B*), with no differences in paired pulse ratios (Extended Data Fig. 4-1*G-H*). Taken together, our data indicate that removal of Ephexin5 in AS mice functionally restores excitatory synapses in the CA1 to levels similar to those observed in WT animals.

**Figure. 4.**
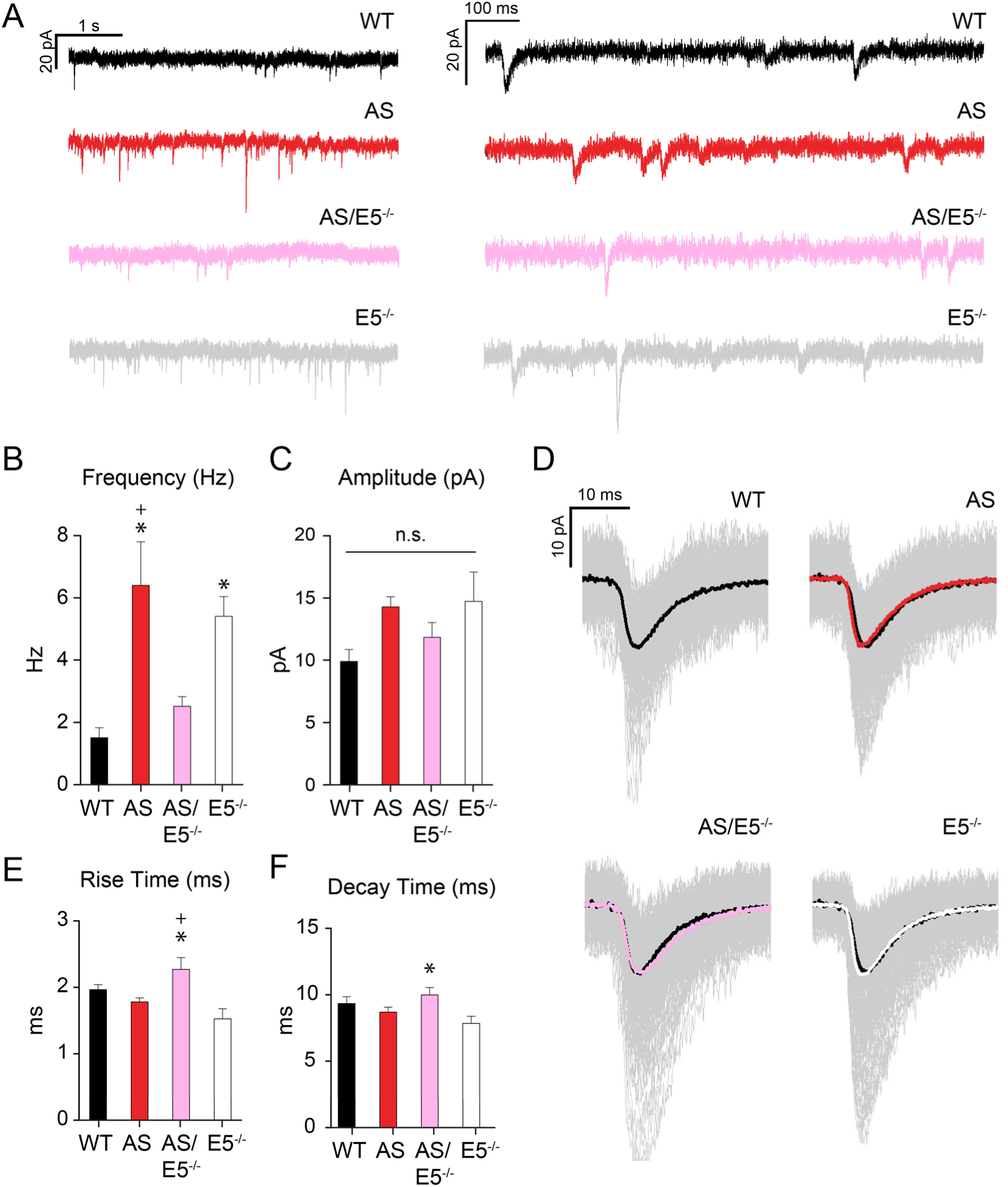
mEPSC frequency, but not amplitude, is altered in the AS CA1 region, and corrected with removal of Ephexin5. **(A)** Representative traces from voltage-clamp recording of each genotype are shown in two different time scales. **(B)** mEPSC frequency recorded from CA1 pyramidal cells represented in Hz. Data are presented as mean ± SEM. \**p*<0.05 compared to WT and ^+^p<0.05 compared to AS/E5^-/-^ (one-way ANOVA) with *post-hoc* Tukey’s multiple comparisons test. **(C)** mEPSC amplitude recorded from CA1 pyramidal cells represented in pA. Data are presented as mean ± SEM. Statistically significant difference between samples was not observed (one-way ANOVA) with *post-hoc* Tukey’s multiple comparisons test. **(D)** Average waveform over individual waveforms for each genotype. WT is shown in black in the other genotype traces for comparison. **(E)** Average rise time for individual cells is shown. Data are presented as mean ± SEM. \**p*<0.05 compared to E5^-/-^, ^+^*p*<0.05 compared to AS (one-way ANOVA) with *post-hoc* Tukey’s multiple comparisons test. **(F)** Average decay times for individual cells are shown. Data are presented as mean ± SEM. \**p*<0.05 compared to E5^-/-^ (one-way ANOVA) with *post-hoc* Tukey’s multiple comparisons test. See also **Extended Data Fig. 4-1**. Sample size (n), degrees of freedom, and exact *p* values are reported in **Table 2** and **Table 3**.

### Removal of UBE3A leads to an activity-dependent increase in spine formation that requires Ephexin5

Why does loss of UBE3A lead to an increase in spine density in an Ephexin5-dependent manner? To address this mechanistic question, we used primary neuronal cultures. Primary hippocampal neurons were co-transfected with plasmids expressing a UBE3A shRNA (or scrambled shRNA) and plasmids expressing GFP. Neurons were then fixed three days post transfection for confocal imaging. We confirmed the knockdown of UBE3A using immunocytochemistry (Fig. 5A). Consistent with previous publications, acute cell-autonomous knockdown of UBE3A did not alter baseline spine density within three days post transfection (Greer et al., 2010).

**Figure. 5.**
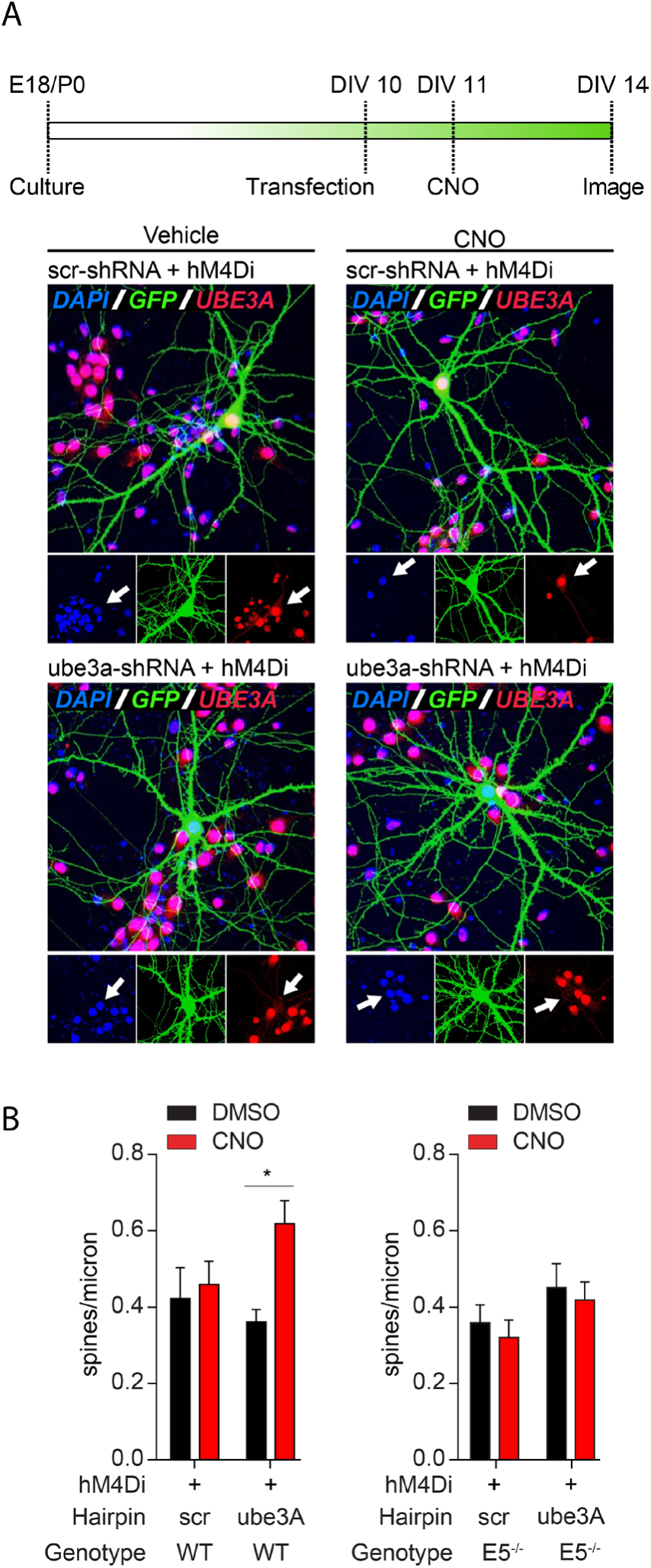
Knockdown of UBE3A leads to an activity dependent increase in dendritic spine formation that requires Ephexin5. **(A)** Hippocampal neuronal cultures were transfected at DIV 10 with plasmids containing GFP, an shRNA hairpin, and hM4Di, and treated with CNO or vehicle from DIV 11-14. Representative neurons at DIV 14 are shown, with white arrows indicating the soma of the transfected cells. Cells were stained for UBE3A (red) and GFP (green). Nuclei were labeled with Hoescht. **(B)** For each treatment group, spine density is shown as spines per micron across the entire transfected neuron. Data are presented as mean ± SEM (n = 3 for all genotypes). \**p*<0.05 (two-way ANOVA) within genotype with *post-hoc* Sidak’s multiple comparisons test. Exact *p* values are reported **Table 2** and **Table 3**.

Previous reports identified decreased synapse density in the CA3 of AS mice (Greer et al., 2010), which could lead to reduced input onto CA1 neurons. Given the cell autonomous nature of our *in vitro* UBE3A knockdown, the transfected cells should in theory receive normal input from surrounding WT neurons. To more closely mimic an *in vivo* AS setting, we cultured hippocampal neurons from WT mice and co-transfected plasmids expressing a UBE3A shRNA or Control shRNA along with a plasmid expressing the G_i_-associated DREADD (hM4Di) (Armbruster et al., 2007) and a plasmid expressing GFP. This approach gave us the ability to cell-autonomously remove UBE3A expression and use the DREADD ligand clozapine N-oxide (CNO) to reduce electrical activity within these neurons as a proxy for reduced excitatory inputs. In transfected neurons, knockdown of UBE3A expression combined with CNO treatment lead to a significant increase in spine density (Fig. 5B). Control shRNA treated neurons were unaffected by CNO treatment. Consistent with our hypothesized role for Ephexin5 in UBE3A-dependent spine regulation, the same manipulation in Ephexin5 knockout primary hippocampal neurons did not lead to increased spine density. From these results, we conclude that hippocampal neurons devoid of UBE3A display aberrant activity-dependent regulation of dendritic spine density, which requires the presence of Ephexin5. Although many more questions remain, we believe that these data provide a clue into how loss of UBE3A and dysregulation of at least one substrate may change communication within hippocampal circuits.

## DISCUSSION

In the present study, we identify Ephexin5 as a substrate of UBE3A. Removing Ephexin5 from AS mice restores the cellular and electrophysiological deficits observed in CA1 of AS mice and rescues performance in hippocampus-dependent learning and memory tasks to WT levels. Phenotypes on various strain backgrounds have been shown to exhibit different results for a given task (Born et al., 2017; Huang et al., 2013a; Sittig et al., 2016), but our results indicate a general rescue of hippocampal-dependent phenotypes but not hippocampal-independent phenotypes in both strains. Furthermore, the learning and memory deficits were recapitulated in animals in which UBE3A was conditionally removed in a subset of excitatory neurons located primarily in the hippocampus. These conditional knockout animals were bred on a C57Bl/6J background and clearly exhibited a deficit in novel place preference, which was also rescued by removal of Ephexin5. Our findings support a role for UBE3A in suppressing Ephexin5 expression in adulthood (or post-development) to allow for normal development of excitatory synapses in the hippocampus.

### Elevation of dendritic spines and synaptic activity in CA1 of adult AS mice

Our study highlights, in adult AS mice, a previously undescribed elevation in hippocampal dendritic spine density and excitatory transmission. Our data suggest this may come from abnormally elevated Ephexin5 expression which leads to aberrant activity-dependent spine plasticity. Although Ephexin5 negatively regulates hippocampal spine formation early in development (Margolis et al., 2010), recent work has indicated that Ephexin5 is also required for regulating activity-dependent spine formation in developing neurons (Hamilton et al., 2017). In AS, it is possible that increased expression of Ephexin5 outside of this critical synaptogenic window could be responsible for an abnormal increase in spine formation in CA1 that is activity-related. The increase in decay time in the AS/E5^-/-^ animals hints at a possible increase in GluR2-containing AMPA receptors (Geiger et al., 1995), which are typically upregulated during development (Kumar et al., 2002) and correlate with increased spine maturity and stability (Suresh and Dunaevsky, 2017). Thus, removing Ephexin5 may be preventing the aberrant activity-related formation of new, immature spines in AS mice.

Although our findings *in vitro* predict that lowered synaptic input into the CA1 neuron *in vivo* would lead to higher spine density in the CA1 of AS mice, the precise pattern of activity *in vivo* causing deficits in CA1 remains unknown. However, previous reports have identified imbalances in excitatory/inhibitory transmission in AS mice, which could be a major contributing factor to the Ephexin5-dependent spine deficits we observed in these mice (Wallace et al., 2012). Based on our data, UBE3A may control activity-dependent spine formation, keeping spine number in check. UBE3A has predominantly been shown to promote synapse development whereby its removal leads to fewer synapses. Whether these changes are relevant to activity-dependent, UBE3A-mediated control of spines is not known. Moreover, while our data appear unexpected, UBE3A overexpression has been reported to inhibit synapse formation in cortical neurons (Smith et al., 2011). Taken together, UBE3A’s role in activity-dependent and activity-independent brain development and function may be more nuanced than previously appreciated.

### Ephexin5 as a drug target

Ephexin5 is a promising candidate to target in future efforts to treat AS. We previously reported that modulation of Ephexin5 can ameliorate learning and memory deficits in Alzheimer’s disease (AD) (Cook et al., 2019; Sell et al., 2017). Interestingly, a recent study links loss of UBE3A in AD to elevation of Ephexin5 and disease progression (Olabarria et al., 2019). This study also supports Ephexin5 as a substrate of UBE3A in pathological conditions. Furthermore, the expression profile of Ephexin5 (high in early development, low in late development/adulthood and enriched in adult hippocampus) suggests that attempts to reduce its expression are unlikely to have off-target side effects. The lack of behavioral deficits in the Ephexin5 mice also supports the case for exploring Ephexin5 as a viable therapeutic target.

### UBE3A-substrate hypothesis

Our findings provide direct evidence in support of a ‘UBE3A-substrate hypothesis’ of cognitive dysfunction in AS and suggest that modulating these UBE3A-directed substrates can improve UBE3A-related cognitive phenotypes. The specificity of phenotypes that were rescued by removal of Ephexin5 in both the Ube3A^loxp-m+/Cre+^ and UBE3A^m-/p+^ models highlights the importance of identifying additional substrates that could be critical for region-specific UBE3A phenotypes. Notably, removing Ephexin5 did not rescue motor and other non-hippocampus dependent phenotypes. Furthermore, given the strong temporal regulation of Ephexin5 expression, the timing of substrate-directed interventions will likely be important to consider when thinking about distinct AS-related behaviors (Silva-Santos et al., 2015). In addition to AS research, such efforts to better understand UBE3A substrates may also be relevant for autistic spectrum disorders (ASD), as duplication of the 15q11-13 chromosomal region encompassing the *UBE3A* locus is one of the few characterized persistent cytogenetic abnormalities associated with ASDs, occurring in >1-2% of all ASD cases (Cook et al., 1997; Glessner et al., 2009).

---

**Extended Data Fig. 1-1.**
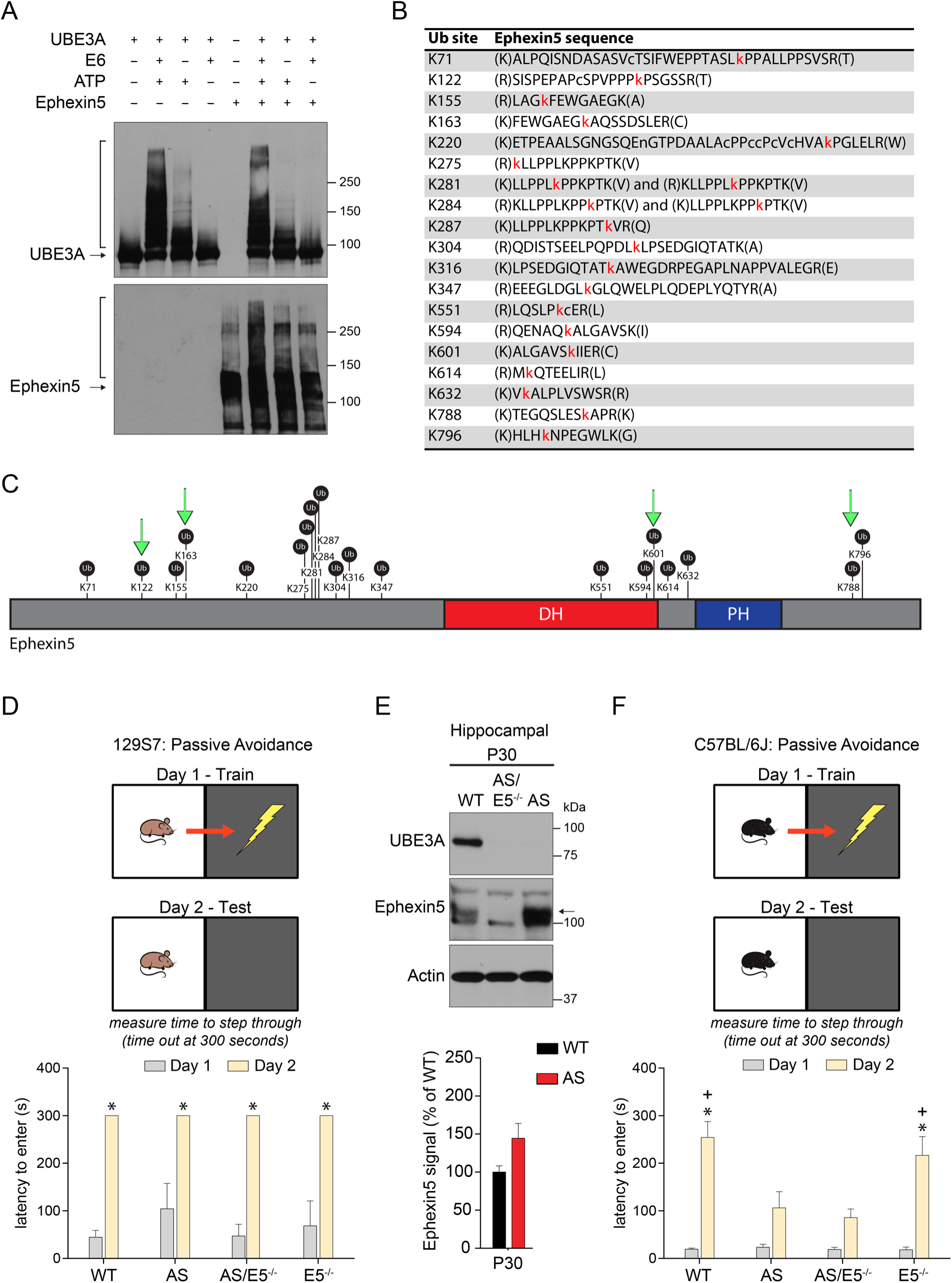
Ephexin5 is ubiquitylated by UBE3A in vitro, has elevated expression in P30 AS hippocampus, and does not contribute to passive avoidance phenotypes, related to Figure 1. **(A)** *In vitro* ubiquitylation assays were performed using purified Ephexin5 protein, with different purified components (UBE3A, ATP, E6, or Ephexin5) of the *in vitro* UBE3A ubiquitylation assay as indicated. The arrows indicate the unmodified protein and the bracket indicates the smearing due to ubiquitylation. Samples were prepared and applied to SDS-PAGE for immunoblot analysis using indicated antibodies. **(B)** Ubiquitylated lysines (red K) identified by tandem mass spectrometry analysis of tryptic peptides from Ephexin 5 after *in vitro* UBE3A ubiquitylation. Green arrows indicate identified sites using reactions in the absence of the enhancing protein E6. **(C)** Diagram of sites of ubiquitylation and domains of Ephexin5. Black circles indicate a site of ubiquitylation at the specified lysine residue. DH = Dbl-homology (Rho-GEF) domain, PH = Pleckstrin homology domain. **(D)** Latency for 129S7 mice to enter shock arena on day 1 and day 2. Data are presented as mean ± SEM. \**p*<0.05 (two-way ANOVA) compared to Day 1 within genotype with *post-hoc* Bonferroni multiple comparisons test. **(E)** Ephexin5 expression levels are elevated in AS mouse hippocampus at P30 and removed in the AS/E5^-/-^. Quantification of Ephexin5 signal is normalized to Actin signal and compared to WT. Arrow head indicates Ephexin5 band. Data are presented as mean ± SEM (n = 6 for WT, n = 7 for AS). **(F)** Latency for C57Bl/6J mice to enter shock arena on day 1 and day 2. Data are presented as mean ± SEM. \**p*<0.05 compared to Day 1 within genotype and ^+^*p*<0.05 compared to Day 2 AS and AS/E5^-/-^ (two-way ANOVA) with *post-hoc* Bonferroni multiple comparisons test. Sample size for behavior (n), degrees of freedom, and exact *p* values are reported in **Table 2** and **Table 3**.

**Extended Data Fig. 2-1.**
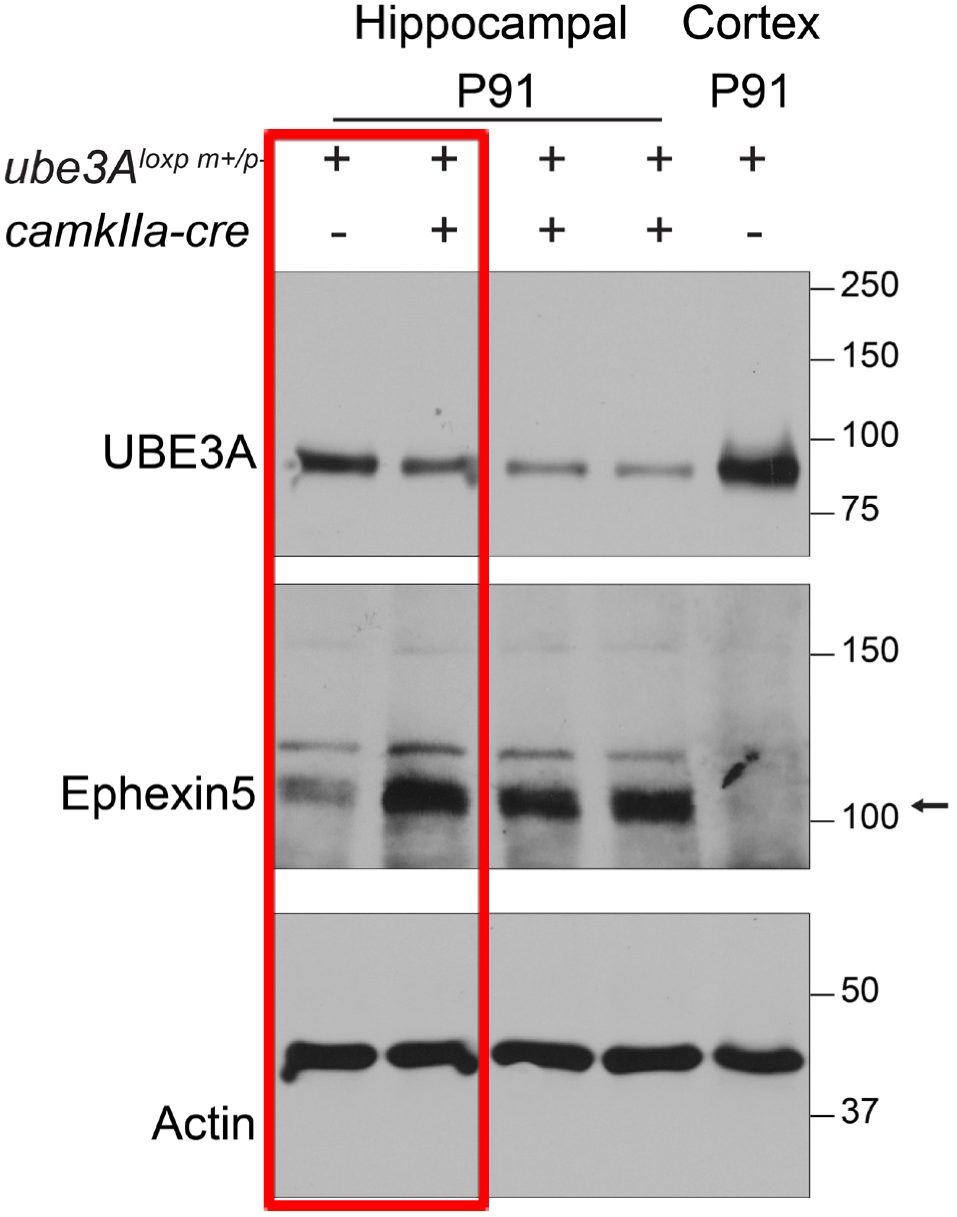
Ephexin5 expression levels are elevated in hippocampus from *ube3A^loxp+^/CamkIIa-cre^+^* mice and not present in the cortex, related to Figure 2. Red box indicates lanes shown in Figure 2B. Additional lanes show increased Ephexin5 expression in hippocampi from *ube3A^loxp+^/camkIIa-cre^+^* mice. Note that Ephexin5 expression is undetectable in cortical samples. Compare lane 5 to lane 1.

**Extended Data Fig. 3-1.**
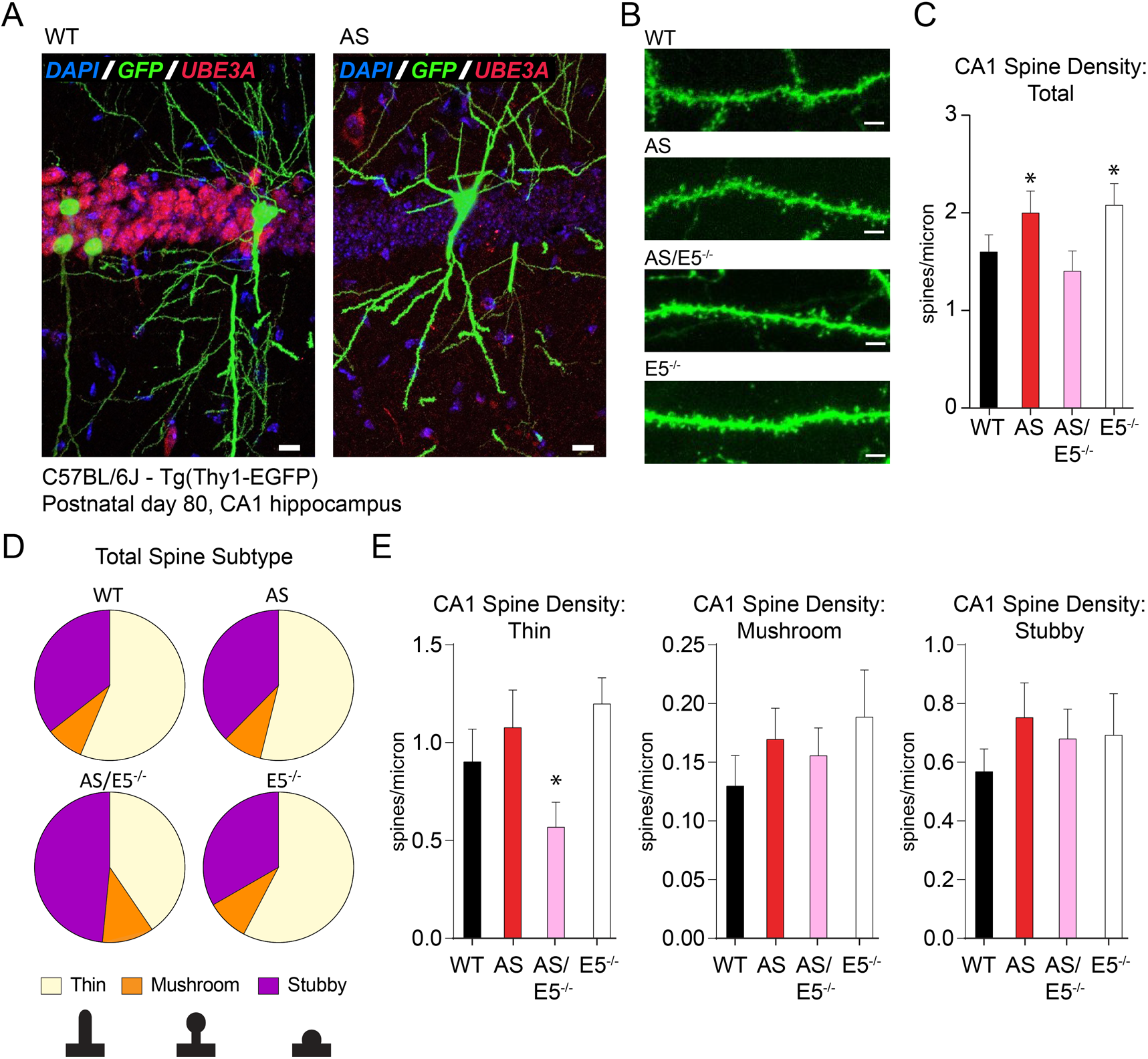
Spine density is altered in the AS CA1 region, and corrected with removal of Ephexin5. **(A)** 11 week old WT and AS Thy1-EGFP-positive tissue was stained for UBE3A (red) and GFP (green), with DAPI (blue) for nuclei labeling. UBE3A is present in CA1 in the WT, but this staining is lost in the AS animal. **(B)** Example dendritic segments from WT, AS, AS/E5^-/-^ and E5^-/-^ CA1. Taken at 63x, scale bar 2 μm. **(C)** Total spine density in CA1. **(D)** Proportion of spine subtypes within WT, AS, AS/E5^-/-^, and E5^-/-^ CA1 neurons. **(E)** Spine density across the CA1 neurons for thin, mushroom, and stubby morphologies. Data are presented as mean ± SEM (n = 3 for all genotypes). \**p*<0.05 (one-way ANOVA) compared to WT and AS/E5^-/-^ with *post-hoc* Tukey’s multiple comparisons test. \**p*<0.05 (two-way ANOVA) compared to AS and E5^-/-^ with *post-hoc* Sidak’s multiple comparisons test for panel. Degrees of freedom, and exact *p* values are reported in **Table 2** and **Table 3**.

**Extended Data 4-1.**
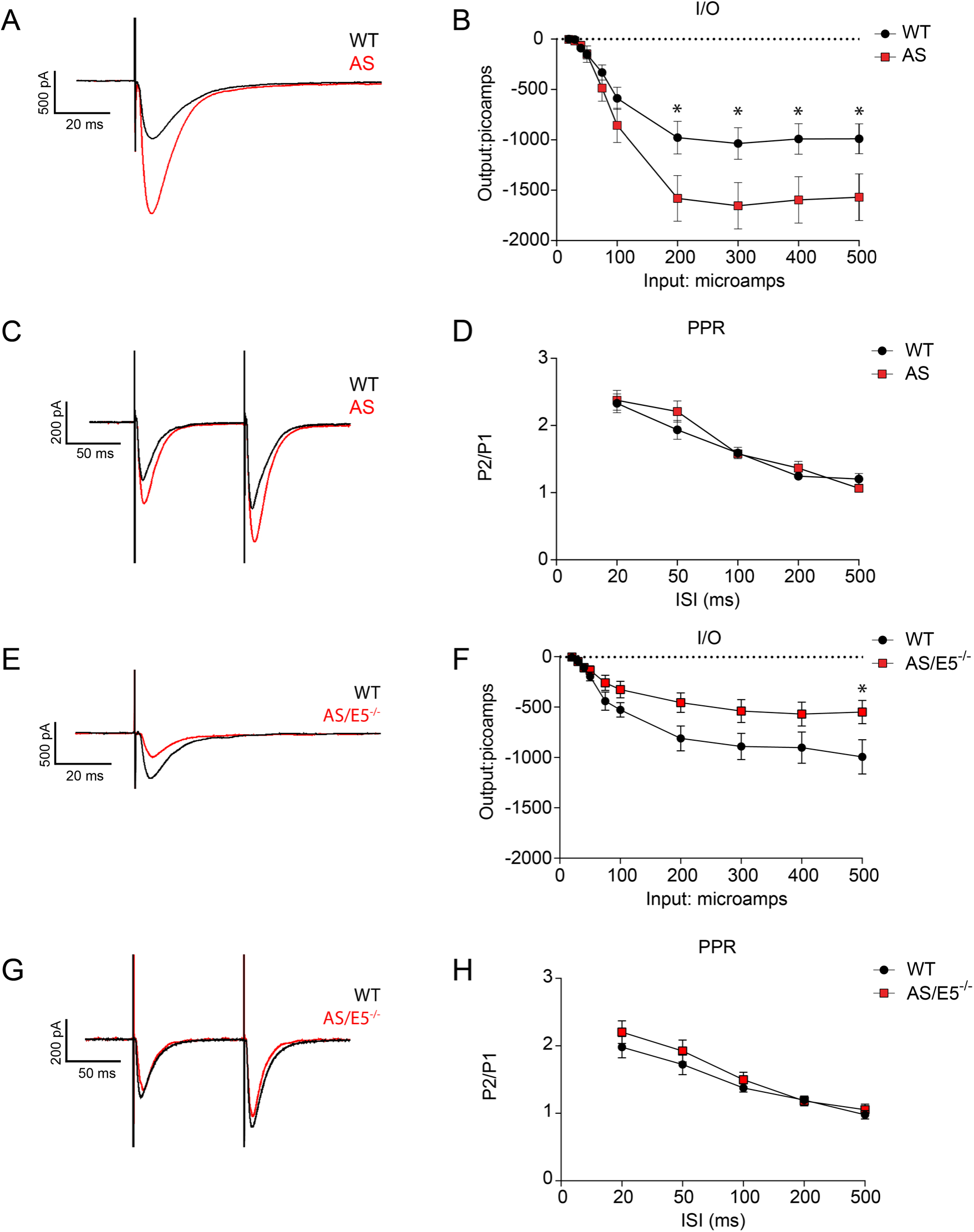
Electrophysiological properties of CA1 pyramidal neurons in WT, AS, and AS/E5^-/-^ hippocampus, related to figure 4. **(A)** Example traces showing evoked potentials in the WT and AS CA1 region after stimulation of the Schaffer collaterals. **(B)** Output in picoamps for WT and AS shown for varying input stimulus intensity. Data are presented as mean ± SEM. One-way ANOVA, *post-hoc* Tukey’s multiple comparisons test. Two-way ANOVA, *post-hoc* Bonferroni multiple comparisons test. \**p*<0.05 compared to WT. **(C)** Example traces showing the evoked potentials in both WT and AS CA1 region after introduction of two stimuli to determine paired pulse facilitation. **(D)** Ratio of second potential over the first potential is shown for varying interstimulus intervals in WT and AS cells. Data are presented as mean ± SEM. Statistically significant difference between samples was not observed (two-way ANOVA) with *post-hoc* Bonferroni multiple comparisons test. **(E)** Example traces showing evoked potentials in the WT and AS/E5^-/-^ CA1 region after stimulation of the Schaffer collaterals. **(F)** Output in picoamps for WT and AS/E5^-/-^ shown for varying input stimulus intensity. Data are presented as mean ± SEM. One-way ANOVA, *post-hoc* Tukey’s multiple comparisons test. Two-way ANOVA, *post-hoc* Bonferroni multiple comparisons test. \**p*<0.05 compared to WT. **(G)** Example traces showing the evoked potentials in both WT and AS/E5^-/-^ CA1 region after introduction of two stimuli to determine paired pulse facilitation. **(H)** Ratio of second potential over the first potential is shown for varying interstimulus intervals in WT and AS/E5^-/-^ cells. Data are presented as mean ± SEM. Statistically significant difference between samples was not observed (two-way ANOVA) with *post-hoc* Bonferroni multiple comparisons test. Sample size (n), degrees of freedom, and exact *p* values are reported in **Table 2** and **Table 3**.

